# Targeted analysis of chondrocyte central metabolites in response to cyclical compression and shear deformations

**DOI:** 10.1101/2025.08.04.668582

**Authors:** Erik P. Myers, Aurora Gallagher, Avery Welfley, Samuel Battles, Aidan Gregory, Priyanka Brahmachary, Mark Greenwood, Ronald K. June

## Abstract

Osteoarthritis results in deterioration of articular cartilage, the soft tissue covering articulating surfaces of bones in joints like the knee and hip. Cyclical mechanical stimulation of articular cartilage results in synthesis of cartilage matrix, suggesting that therapeutic mechanical stimulation might be beneficial for cartilage repair in osteoarthritis. Several prior studies identify ion channels and cytoskeletal molecules as components of chondrocyte mechanotransduction. The pathways of central metabolism, glycolysis, the pentose phosphate pathway, and the tricarboxylic acid cycle are necessary for producing non-essential amino acids that are needed for synthesizing matrix proteins for cartilage repair. However, it is currently unknown if and how levels of central metabolites change with applied mechanical stimulation of chondrocytes. Here, we show that applied cyclical shear and compressive deformations drive changes in multiple central metabolites in primary chondrocytes. These results add to a rich picture of chondrocyte mechanotransduction, including calcium signaling, ion channels, integrins, and cytoskeletal components. By finding compression- and shear-induced changes in central metabolites, these data support the potential for therapeutic mechanotransduction toward cartilage repair. Future studies may build on these results to understand the relationships between mechanical stimulation and chondrocyte central metabolism.

## Introduction

Articular cartilage is the soft tissue lining diarthrodial joints like the knee and hip. This tissue contains a single cell type—the articular chondrocyte—and is subjected to cyclical loads during daily activities such as walking.These loads cause tissue-level deformations that result in mechanical stimulation of chondrocytes. Tissue-level deformations include tension, compression, and shear-based on *in vivo* measurements (Chan et al. 2016).

A key feature of the most common joint disease, osteoarthritis (OA), is the deterioration of articular cartilage (Tonge et al. 2014).This deterioration results in alterations of both the extracellular and pericellular cartilage matrices, with concomitant changes in various cartilage mechanical properties. For example, after joint injury, there are significant decreases in the indentation modulus of the mouse pericellular matrix after just 3 days, with a reduction of ∼50% after 1 month (Chery et al. 2020). In contrast, exercise typically results in improved cartilage health. Patients with a history of marathon running had ∼25% thicker cartilage than age-matched controls (Mosher et al. 2010).

Early observations show that cyclical mechanical stimulation of cartilage and chondrocytes results in production of matrix molecules (Mauck et al. 2006; Sah et al. 1989) through chondrocyte mechanotransduction. This shows that mechanical stimulation can drive therapeutic production of matrix for cartilage repair. While several ion channels (Ely et al. 2025) and various cytoskeletal components (Haudenschild et al. 2011; Ohashi et al. 2006) have been implicated in chondrocyte mechanotransduction, many questions remain regarding the specific cascade of signals that results in increased matrix production.

Central metabolism (*i.e*. energy metabolism) is a network of enzymes and small-molecule intermediates called metabolites used by cells to harvest carbon and energy to support cellular function. Substrates include glucose and glutamine, which can be metabolized to produce non-essential amino acids. Functioning central metabolism, including both glycolysis and cellular respiration, involves oxygen consumption and is essential for producing the non-essential amino acids required for cartilage regeneration. For example, non-essential amino acids make up 83.1, 71.4, and 74.5% of the primary sequences for the key cartilage matrix proteins encoded by COL2A1, COL6A1, and ACAN (types II and VI collagen and aggrecan).

We hypothesized that mechanical stimulation would alter the production of central metabolites in articular chondrocytes. Toward this goal, we harvested articular chondrocytes, expanded them in monolayer culture, and then encapsulated them in physiologically stiff agarose (Jutila et al. 2015; Zignego et al. 2014). Embedded chondrocytes were then stimulated with cyclical shear and compressive strains at physiological levels. Immediately after mechanical stimulation, metabolites were extracted. Quantification of central metabolites was performed using targeted metabolomic profiling. We found changes in levels of several metabolites, demonstrating that mechanical stimulation affects chondrocyte central metabolism. These results can inform future studies on how mechanical stimuli influence chondrocyte metabolism for cartilage maintenance and repair strategies.

## Methods

### Chondrocyte Harvest and Culture

To assess central metabolism in chondrocyte mechanotransduction, chondrocytes were isolated from cartilage using established methods (Zignego et al. 2015; Zignego et al. 2014). Primary Human Chondrocytes (PHCs) were obtained from discarded tissue from Stage-IV osteoarthritis patients (n=5 male and n=5 female) undergoing total joint replacement under IRB approval. Bovine chondrocytes (BCs, n=5) were harvested from knee joints of 18–22-month-old cattle obtained from a local abattoir. Articular cartilage was digested with Type I Collagenase (2 mg/mL in DMEM with penicillin and streptomycin, Gibco, Waltham, MA, USA) for 14 h at 37 ºC. Isolated chondrocytes were cultured in Dulbecco’s Modified Eagle’s medium (DMEM, Gibco, Waltham, MA, USA) supplemented with Fetal Bovine Serum (FBS, 10% v/v, Bio-Techne, Minneapolis, MN, USA), penicillin (10,000 I.U./mL), and streptomycin (10,000 µg/mL, Sigma, St. Louis, MO, USA) in 5% CO2 at 37 ºC. First passage cells were used for experiments.

To provide a three-dimensional environment in the range of physiological stiffness, chondrocytes were encapsulated in high-stiffness agarose (Jutila et al. 2015). After trypsinization, cells were washed with 1X PBS and resuspended in DMEM (without phenol red) and mixed with low-melting-temperature agarose (Type VII-A Sigma Aldrich) at a final concentration of 4.5% wt/vol with ∼500,000 cells/gel and allowed to solidify for 10 min. Hydrogels were cast in a cylindrical aluminum mold to have a nominal diameter of 0.5” and a height of 0.25” following a previously established protocol (Jutila et al. 2015), which results in chondrocyte viability >90% (Zignego et al. 2014). Agarose gels were placed in 24-well tissue culture plates (Falcon) with 2 mL custom DMEM media containing 4.5g/L glucose, 2 mM L-glutamine,110mg/L sodium pyruvate, 10% FBS, and 1% PenStrep and cultured overnight in 5% CO_2_ at 37ºC. Prior to loading, media was replaced with PBS for 1 hr and incubated in 5% CO_2_ at 37ºC.

### Mechanical Stimulation

To evaluate the effects of physiological loading on chondrocyte central metabolism, agarose-embedded chondrocytes were subjected to different mechanical stimuli. Stimuli included applied cyclical shear and compressive strains in a tissue culture environment (Figure 1, Supplemental Material 1). A total of N=10 PHCs (n=5 male and n=5 female) and N=5 BC samples were used.

**Figure 1:**
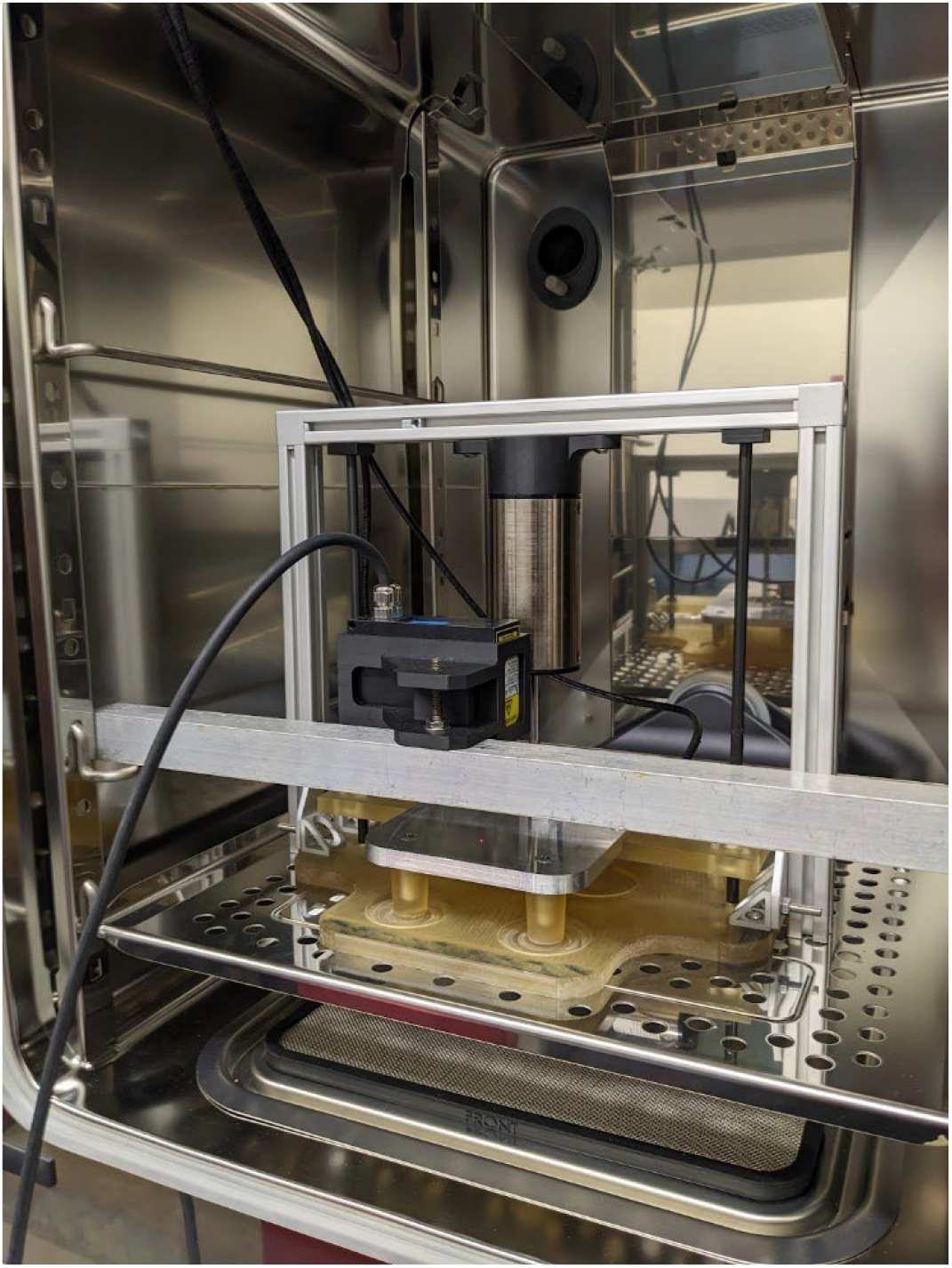
Chondrocyte bioreactor inside a Tri-Gas incubator with laser displacement sensor suspended overhead. During normal experimentation, the sensor is absent from the incubator.

Chondrocyte-embedded agarose constructs from each bovine or human donor were randomized into five experimental groups: unloaded controls (U), 15 minutes of cyclical compressive strain (15C), 15 minutes of cyclical shear strain (15S), 30 minutes of cyclical compressive strain (30C), and 30 minutes of cyclical shear strain (30S). For both compressive and shear stimulation, mean strains were 5% with an amplitude of 2.5%. Each experiment was performed at 1.1Hz with 5% preload applied initially for 15 minutes to allow for stress relaxation. Mechanical stimulation of samples from each donor was completed over the course of a single day. In instances where multiple donors were available on the same day, simultaneous loading for each group across all available donors was performed.

For each loading group, the hydrogel was removed from PBS, placed within the polysulfone sample cup, and resubmerged in fresh PBS to prevent drying during loading (Figure 1). The bioreactor was then placed inside the Tri-GAS incubator and powered on to provide mechanical stimulation within a tissue culture environment. In the case of an uneven number of hydrogels, a cell-free agarose sample was used to balance the load from the hydrogels on the plunger plate.

Immediately after mechanical stimulation, hydrogels were removed from the bioreactor and bisected using a sterile scalpel. One-half of the sample was used for oxygen measurement and the other half was used for targeted metabolomic analysis.

### Oxygen Measurement and Metabolite Extraction

An oxygen measurement system (OX-10, Unisense, Aarhus, Denmark) was used to quantify oxygen tension immediately after mechanical stimulation. Probes were calibrated to 0% O_2_ tension using a nitrogen sparged water bath and 100% O_2_ tension using an atmospheric air sparged water bath. 2-point calibration curves were then used to quantify oxygen saturation readings from each sample bisect. The oxygen probe was inserted into the hydrogel and oxygen readings were measured for 1 minute, with the minimum oxygen tension during the measurement recorded for each sample (Figure 2).

**Figure 2:**
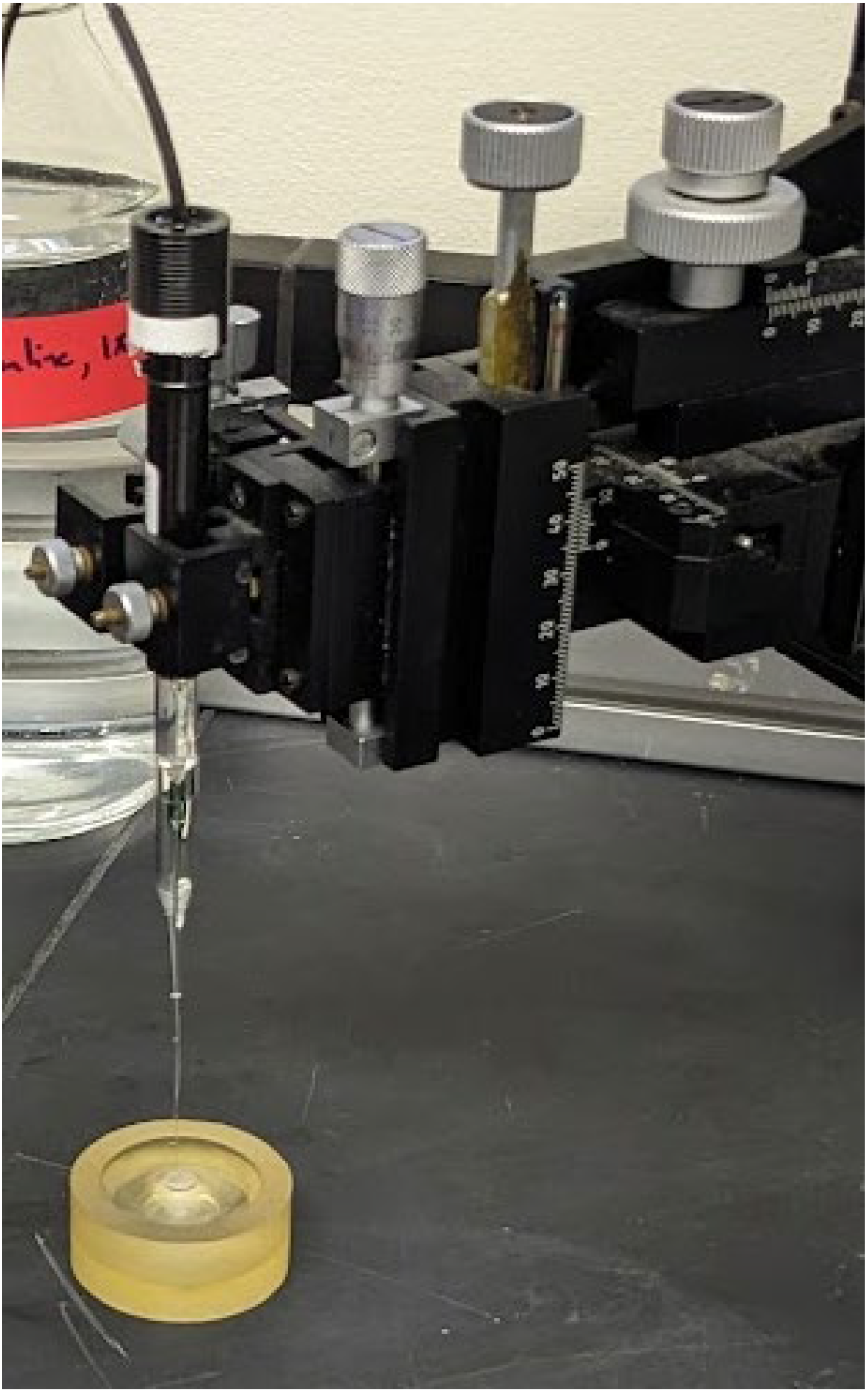
Oxygen probe needle embedded within an agarose hydrogel. The probe was retracted after each reading and used to collect a pre and post reading atmospheric baseline.

The remaining half of the hydrogel was immediately wrapped in autoclaved aluminum foil and flash-frozen in liquid nitrogen to preserve metabolite levels after mechanical stimulation. After freezing for 1-2 minutes, the gel was crushed using a sterile aluminum rod. The pulverized hydrogel was then placed in a 2.5mL Eppendorf tube and allowed to freeze in -80°C overnight. This process was completed for all hydrogels for all experimental groups. After freezing for at least 12 hours, the crushed gels were removed from the freezer and homogenized into a powder using a Cryomill (Verder Scientific, Haan, Germany).

Following homogenization, the powder was suspended in extraction buffer (70:30:1% Methanol:Acetone:Formic acid). Formic acid facilitated metabolite release from the homogenized agarose (Supplemental Material 2). Each hydrogel bisect was extracted with 5mL of extraction buffer at -80°C overnight. The following morning, samples were centrifuged (4,000 rpm, 10 minutes), and the supernatant was evaporated using a vacuum centrifuge at 45°C (Eppendorf, Hamburg, Germany) until only the dried pellet of concentrated metabolites remained. These metabolite pellets were then resuspended in 50µL 50:50 HPLC water:acetonitrile and sent for mass spectrometry analysis.

### Mass spectrometry analysis of mechanically stimulated constructs

After metabolite extraction, levels of central metabolites were quantified using liquid chromatography-mass spectrometry (LC-MS) using an established protocol (Brahmachary et al. 2023) optimized for an Agilent 6538 Q-TOF (C18 Column). Quality control included analysis of blank samples of 50:50 HPLC Water:Acetonitrile every ten samples. For quantification, analytical standards containing 18 key central carbon metabolites were included within the run at concentrations of 1-100 µg/mL. Individual calibration curves were constructed based on the known standard concentration and the measured metabolite intensity (Supplemental Material 3). A pool was created for each loading group and assembled from 5 randomly chosen 10µL samples from each group to aid in fragmentation and identification of each standard metabolite in the sample. After LC-MS analysis, data were converted to concentrations from intensities using the calibration curves before statistical analysis and visualization.

### Statistical Analysis of Metabolites

After removing outliers due to metabolite extraction challenges (Supplemental Material 2), the data from the remaining n=3 male human, n=3 female human, and n=5 bovine donors were combined for a total of 65 samples and analyzed on a per-metabolite basis. Eighteen standard metabolites were used for mass spectrometry calibration; two different forms of NADH as well as HSCoA and oxaloacetate were not detected consistently in the experimental samples, leaving a total of 14 central metabolites for comparison (Supplemental Material 2-3). Metabolites, their pathways within central carbon metabolism, and the significance of mean differences with the unloaded controls were recorded for all 14 detected metabolites (Table 1).

**Table 1:** Individual metabolites, their pathways, and statistical results. ANOVA p- value is the p-value for the repeated measures ANOVA. Significant groups indicate groups significantly different from unloaded controls with arrows indicating increased (↑) or decreased (↓) levels.

| Metabolite | Pathway | ANOVA p-value | Significant Groups vs Unloaded Ctrl |
| --- | --- | --- | --- |
| Fructose / Glucose (Glucose) | Glycolysis | 0.035 | ↓30C, ↓30S |
| Fructose-6-phosphate | Glycolysis | 0.488 |  |
| Phosphoenolpyruvic Acid (PEP) | Glycolysis | 0.641 |  |
| Pyruvic Acid (Pyruvate) | Glycolysis | 0.0046 | ↓30C |
| Lactic Acid (Lactate) | Glycolysis | 0.220 |  |
| 6-p-gluconate | PPP | 0.747 |  |
| Sedoheptulose-7-phosphate | PPP | 0.187 |  |
| Citric Acid (Citrate) | TCA Cycle | 0.615 |  |
| 2-Oxoglutaric Acid (α-ketoglutarate) | TCA Cycle | 0.0984 | |
| Succinic Acid (Succinate) | TCA Cycle | 0.125 | ↑30S |
| Fumaric Acid (Fumarate) | TCA Cycle | 0.358 |  |
| Malic Acid (Malate) | TCA Cycle | 0.0005 | ↓30C, ↓30S |
| Glutamic Acid (Glutamine) | TCA Cycle | 0.211 |  |
| Glutamine | TCA Cycle | 0.0085 | ↓15C, ↓30C, ↓15S, ↓30S |

For each of the detectable 14 metabolites, statistical analysis was performed using Graphpad Prism. Outlier identification was performed using the method of Motulsky and Brown (ROUT in Prism) with a false discovery rate of 1% (Motulsky and Brown 2006). One-way repeated measures ANOVA was performed on log-transformed data to test for differences in mean concentrations between the experimental groups. To control the family-wise error rate, additional pairwise comparisons were performed using Tukey’s HSD. T-tests were used to compare metabolite levels between chondrocytes from male and female donors using the Holm-Sidak correction for multiple testing between experimental groups without assuming sphericity. Metabolites with significant differences are visualized using violin plots. All metabolite data is available in Supplemental Material 4.

## Results

### Oxygen Concentration

Because oxygen is consumed during cellular respiration, oxygen readings were sampled for each hydrogel during experimentation. Statistical analysis of the oxygen readings revealed no significant differences in oxygen tension regardless of loading condition. This may indicate that there was not sufficient time for chondrocytes to utilize measurable oxygen during the short time span (0-30 minutes) of the mechanical stimulation. Oxygen data are available in Supplemental Material 5.

### Targeted Metabolomics

Toward understanding how mechanical stimuli affect chondrocyte central metabolite levels, we performed targeted metabolomic profiling after cyclical shear and compressive stimulation. Five of the fourteen measured metabolites resulted in at least one significant between-group difference. Of these five, two are related to glycolysis (Figure 3) and three are related to the TCA cycle (Figure 4). Four metabolites decreased in quantity in the mechanically stimulated groups, suggesting that mechanical stimulation either decreased production or increased consumption of these metabolites. For the human OA donors, there were no detected differences in any measured metabolite levels between male and female donors (minimum n=3 each). Oxaloacetate, NADH1, HSCoA, and NADH2 were not consistently detected amongst the samples and were not analyzed further, and there were no differences in pentose phosphate metabolites (Figure 5.).

**Figure 3:**
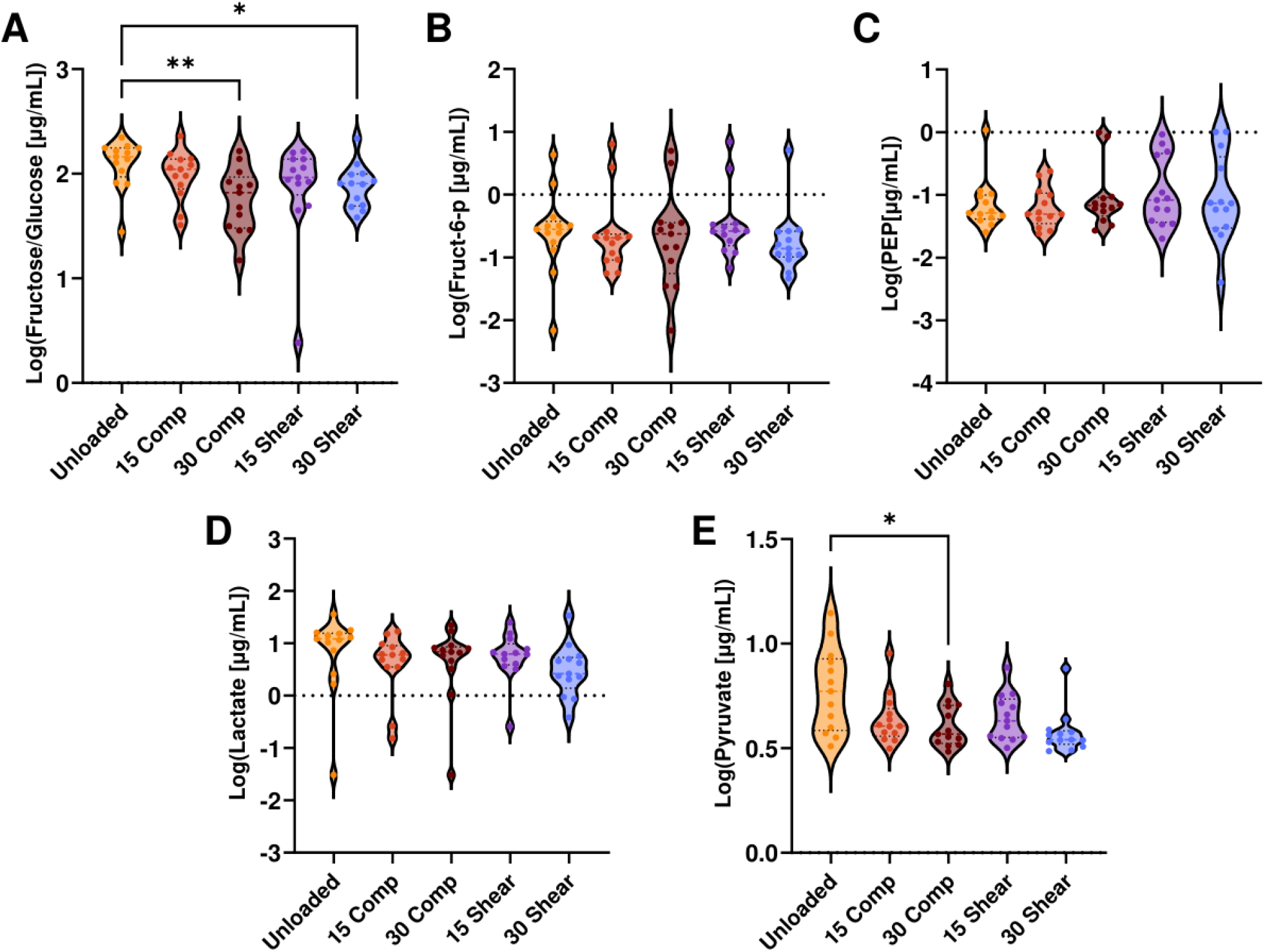
Concentrations of glycolytic metabolites in response to shear and compressive mechanical stimulation. (A) Fructose/Glucose. (B) Fructose-6- phosphage. (C) Phosphoenolpyruvate. (D) Lactate. (E) Pyruvate. * indicates p<0.05 and ** indicates p<0.01 upon Holm-Sidak post hoc test between experimental groups. n=11 per group.

**Figure 4:**
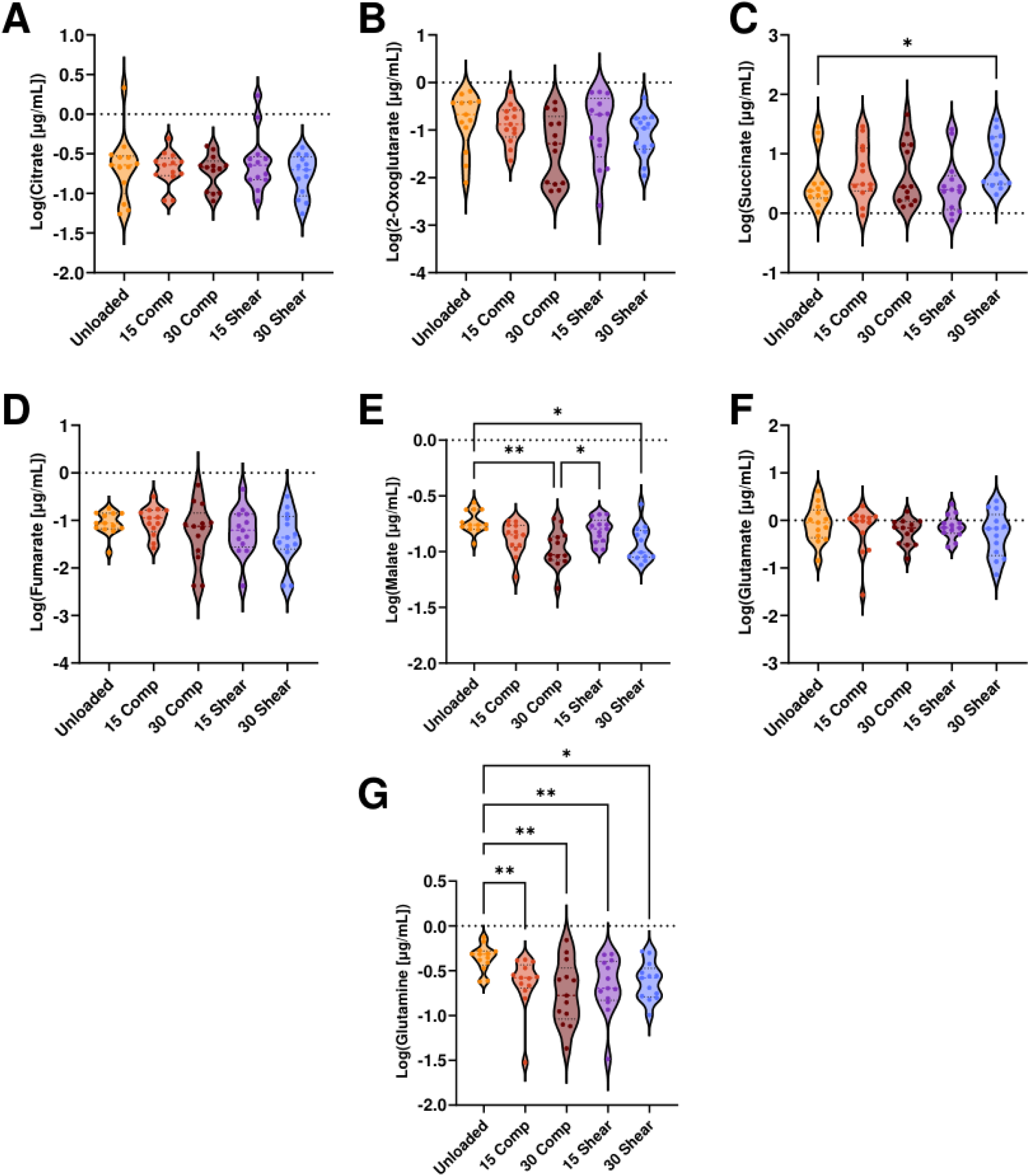
Concentrations of TCA metabolites in response to shear and compressive mechanical stimulation. (A) Citrate. (B) 2-Oxoglytarate. (C) Succinate. (D) Fumarate. Malate. (F) Glutamate. (G) Glutamine. * indicates p<0.05 and ** indicates p<0.01 upon Holm-Sidak post hoc test between experimental groups. n=11 per group.

**Figure 5:**
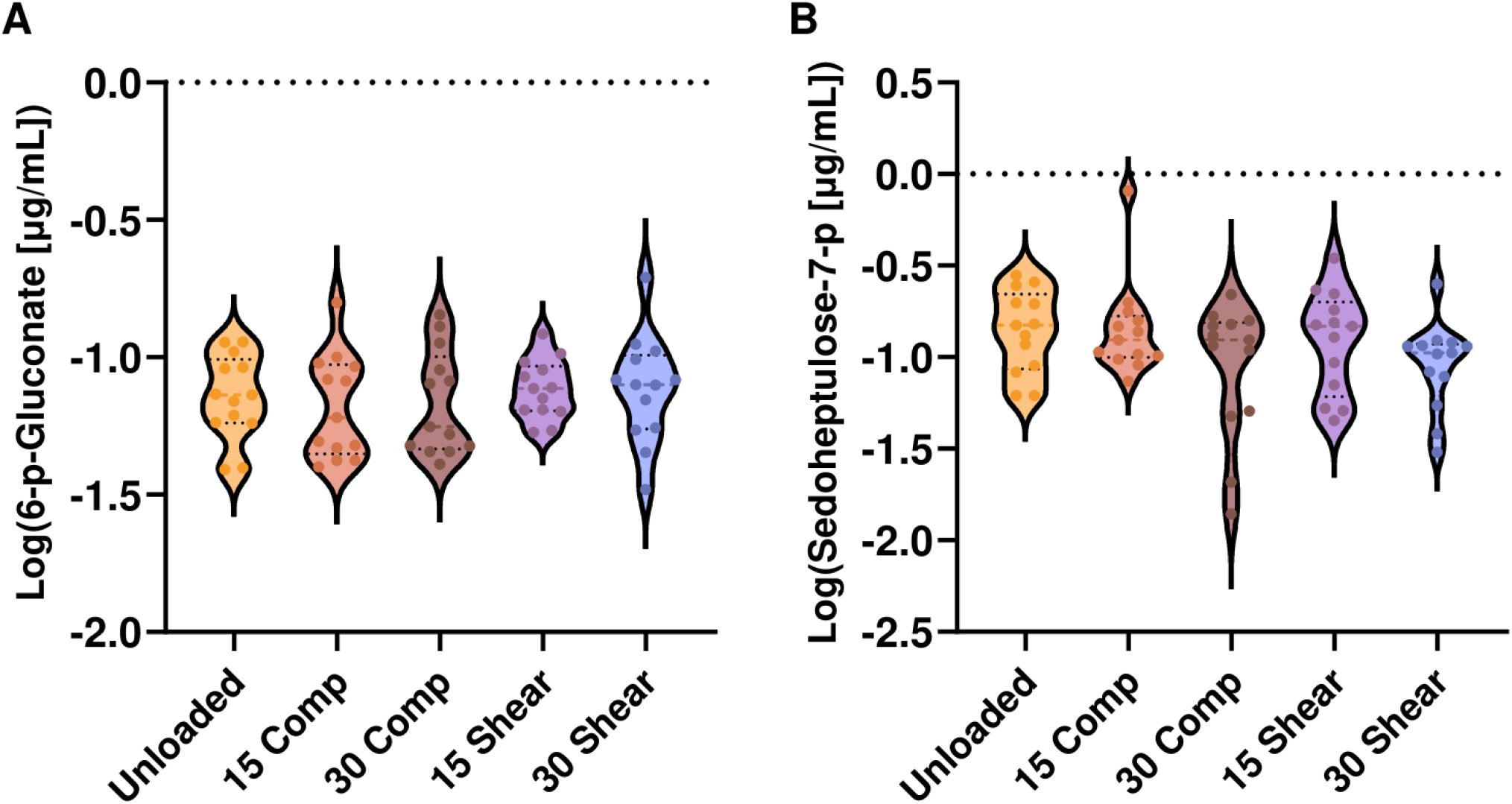
Concentrations of Pentose Phosphate Pathway metabolites in response to shear and compressive mechanical stimulation. (A) 6-phospho-gluconate. (B) sedoheptulose-7-phosphate. No significant differences observed. n=11 per group.

Glucose, the input to glycolysis, decreased after both 30 minutes of cyclical compression and shear compared to unloaded controls (Figure 3A). Glucose concentrations were substantially higher than other metabolites because of the initial culture in media with 4.5g/L glucose. Pyruvate, an end product of glycolysis, decreased in response to 30 minutes of shear and compression stimulation (Figure 3E). In anaerobic respiration, pyruvate can be converted to lactate, but there were no significant differences in mean lactate after mechanical stimulation. These results indicate that there was substantial consumption of glycolytic metabolites in response to both applied shear and compression for chondrocytes.

Within the TCA cycle, succinate increased after 30 minutes of shear stimulation compared to controls (Figure 4C). Malate levels decreased after 30 minutes of both compression and shear stimulation (Figure 4E). Glutamine, a carbon source for the TCA cycle, levels decreased for both 15 and 30 minutes of compression and shear compared to unloaded controls (Figure 4G). These data indicate that cyclical shear and compression induced substantial TCA activity in articular chondrocytes.

## Discussion

This study examined the role of cyclical mechanical stimulation in chondrocyte metabolism using targeted profiling of metabolites from glycolysis, the pentose-phosphate pathway, and the TCA cycle. Mechanical loading altered several key metabolites, revealing that chondrocyte metabolism dynamically responds to cyclical shear and compressive stimuli. However, given the complexity of these larger biochemical networks, changes in metabolite abundance may reflect altered production and/or consumption (Doran et al. 2024).

Glycolytic metabolites (glucose and pyruvate) were decreased in mechanically stimulated samples compared to unloaded controls, suggesting decreased production or increased consumption, or a combination thereof. This may indicate a shift toward a glycolytic, Warburg-like profile often seen in pathological conditions (Jiang et al. 2023) or increased fueling of the TCA cycle to promote production of non-essential amino acids for production of matrix proteins. Glutamine—a primary carbon source for the TCA cycle— was similarly reduced in all mechanically stimulated groups compared to unloaded controls, demonstrating changes in glutamine metabolism because of mechanical stimulation. This reduction in glutamine following mechanical loading suggests enhanced glutaminolysis, where glutamine is converted to glutamate and subsequently to α- ketoglutarate to support increased TCA cycle activity. In parallel, it is possible that glutamate can also be diverted toward proline biosynthesis to support matrix remodeling, together explaining the observed drop in glutamine levels.

Conversely, succinate levels increased after 30 minutes of shear stimulation, which may be linked to downstream redox signaling or mitochondrial stress, as supported by findings in osteoarthritic bone tissues (Shodiev et al. 2024). Collectively, this altered TCA activity suggests that mechanical loading increases energy demand and may shift chondrocyte metabolism toward pathways that support the production of non-essential amino acids for protein and matrix production.

Beyond this system, our findings may have broader implications for cyclical loading in osteoarthritis progression. The effect of mechanotransduction on central metabolism and signaling is increasingly recognized in all joint tissues (Zhao et al. 2020). These results support the emerging evidence that metabolism is not a passive outcome of chondrocyte function, but rather a key response to mechanical signals. In healthy tissue, enhanced mitochondrial metabolism under physiological loading supports ECM synthesis and redox homeostasis (Coleman et al. 2016). In contrast, excessive loading can promote glycolytic upregulation and impaired mitochondrial respiration, leading to an accumulation of metabolic intermediates and reactive oxygen species that activate inflammatory signaling cascades and cytokine expression (Huang et al. 2023; Tan et al. 2022). Thus, these mechanically induced metabolomic changes may serve as both a mechanism of adaptation and a contributor to degeneration, depending on the cellular context and load magnitude, underscoring their potential as therapeutic modulators of cartilage metabolism.

A key limitation of this study is that these targeted metabolomic profiles only provide a static snapshot of metabolite levels at 15-minute time intervals, limiting the interpretability of metabolic flux. Additionally, measuring oxygen consumption immediately after loading may not provide sufficient time for chondrocytes to appreciably consume oxygen within the agarose, and later timepoints may provide additional insights. Future studies should utilize stoichiometric modeling to better understand how mechanical stimulation influences flux of central metabolites and might also incorporate improved temporal resolution with metabolite biosensors in live cell imaging, functional assays, and isotope tracing to better resolve metabolic flux and pathway dynamics. These methods would help quantify nutrient utilization and identify specific biochemical reactions altered by mechanical stimuli. Given the small sample size (n=3 each) of male and female donors, future studies should further explore potential sex differences, hormonal influences, and mitochondrial regulation. These findings could ultimately guide regenerative medicine approaches for OA by identifying optimal mechanical loading parameters that support metabolism-based repair and maintenance of articular cartilage.

## Acknowledgements

This study was funded by research grants from the NIH (NIAMS R01AR073964 and R01AR081489) and NSF (CMMI 1554708).

## Supplemental Material

1. Bioreactor Design
2. Metabolite Extraction and Outlier Analysis (formerly Supp Mat B)
3. Metabolite Calibration Curves (formerly Supp Mat C)
4. Raw Metabolite Data
5. Raw Oxygen Data

