## Supplemental File 1 for "Targeted analysis of chondrocyte central metabolites in response to cyclical compression and shear deformations"

**Supplemental File 1: Bioreactor Design and Drawings**

Seven total design criteria (DC) were established and used to guide the design process of the hydrogel bioreactor:

1. Total bioreactor weight and size should be easily handled by any lab technician and comfortably relocated to within a trigas incubator.
2. Bioreactor design should be corrosion resistant and able to operate within a hot and humid environment for periods exceeding 30 minutes.
3. Bioreactor control should be achievable with a GUI requiring no prior programming experience or knowledge of internal machine code operation.
4. Bioreactor operation should be independent of computer interface in the event of hardware disconnection (headless operation).
5. Bioreactor should retain the ability to accommodate at least six (6) simultaneous samples.
6. Loading profiles should be expanded to include both compressive and shear capabilities.
7. Loading regimes should be easily modifiable on the fly and include modifiable loading conditions including loading time, compressive strain amplitude, shear strain amplitude, loading frequency, preload amplitude, and preload time.

The initial bioreactor design was based loosely on the format of modern 3-d printers like the Ender 3 Pro. In this design, a gantry suspends actuated components above a working volume. Driven by DC #1 and #2, primary design materials were selected to be both 6061 aluminum and polysulfone. These materials are corrosion resistant, relatively easy to machine, are safe to autoclave, and readily available

(McMaster-Carr). CAD work was performed primarily in SolidWorks Design Suite. Post-Processing CAM G-code was produced using Fusion-360 for each sub-assembly component exported from SolidWorks. Milling work was done primarily using a SuperMax-YCM 30 CNC endmill and Fagor 8025 controller. Lathe work was performed using an Enco 1236 metal lathe. All components which were not readily available for purchase were designed and manufactured in-house.

Given DC #5, the base plate was designed to be a flat polysulfone plate with indents (Figure S1).

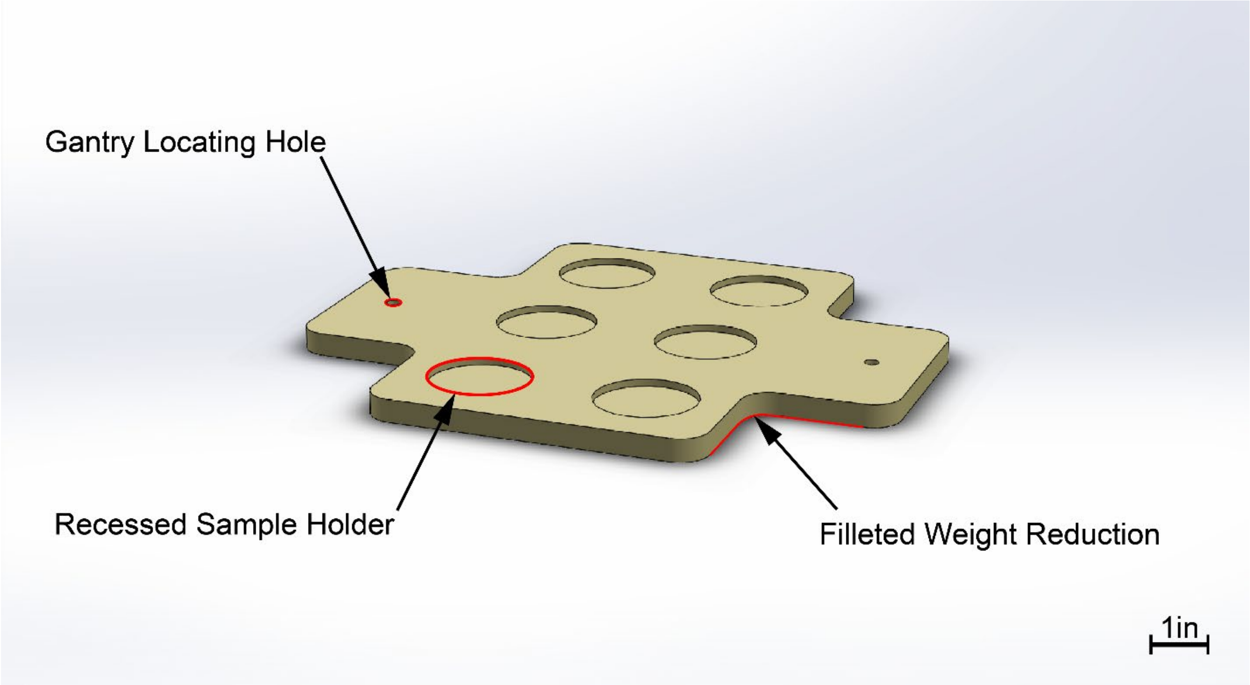


*Figure S1: Base plate made from polysulfone. Six indents are machined away to index the six sample cups. Two locating holes for the gantry rods are drilled. Weight is reduced by removing material in the corners of the baseplate and filleting sharp geometry changes.*

The base plate was made from polysulfone to maximize the reduction of weight over 6061 aluminum, given that the base plate receives support from the laboratory bench or incubator shelf during operation. To hang the gantry and operating motors, a superstructure was constructed from readily available 80/20 20mm anodized 6061 aluminum (McMaster-Carr) (Figure S2).

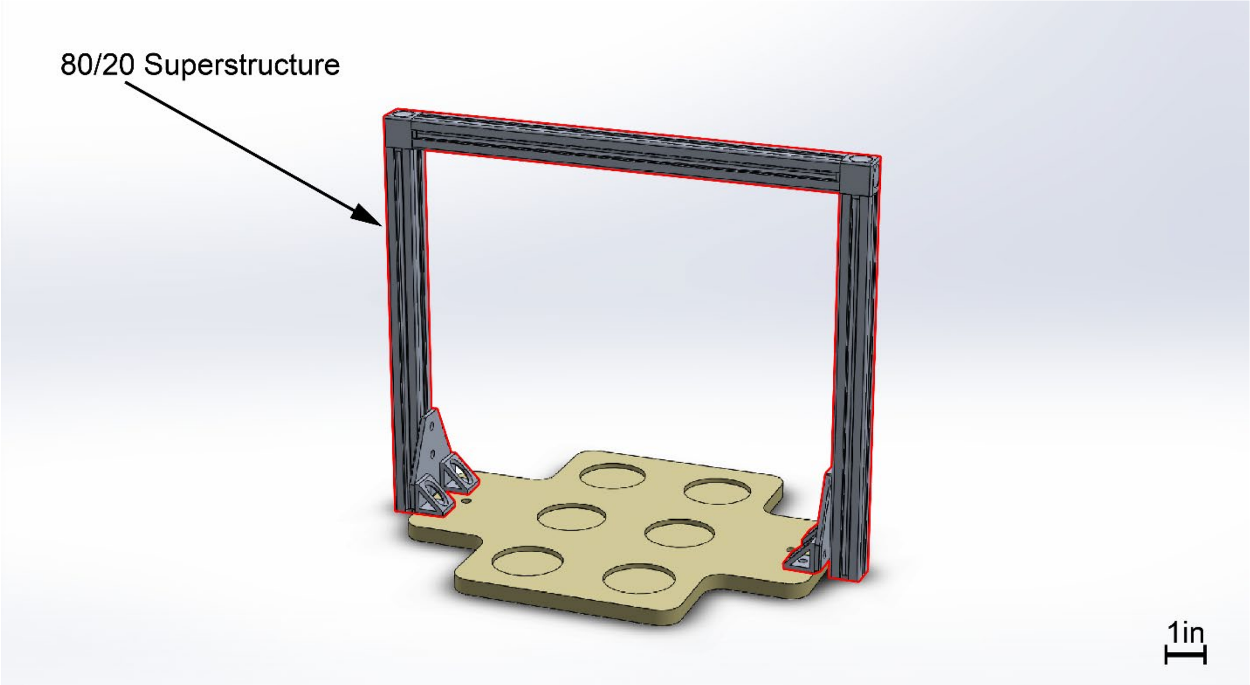


*Figure S2: 80/20 support structure above the base plate. Anodized 20mm railing provides horizontal and vertical connections for interfacing attachments.*

Once the superstructure of the bioreactor was assembled, the articulating components could be incorporated. A horizontal gantry was suspended over the base plate using the pre-drilled gantry locating holes and locating rods (Figure S3).

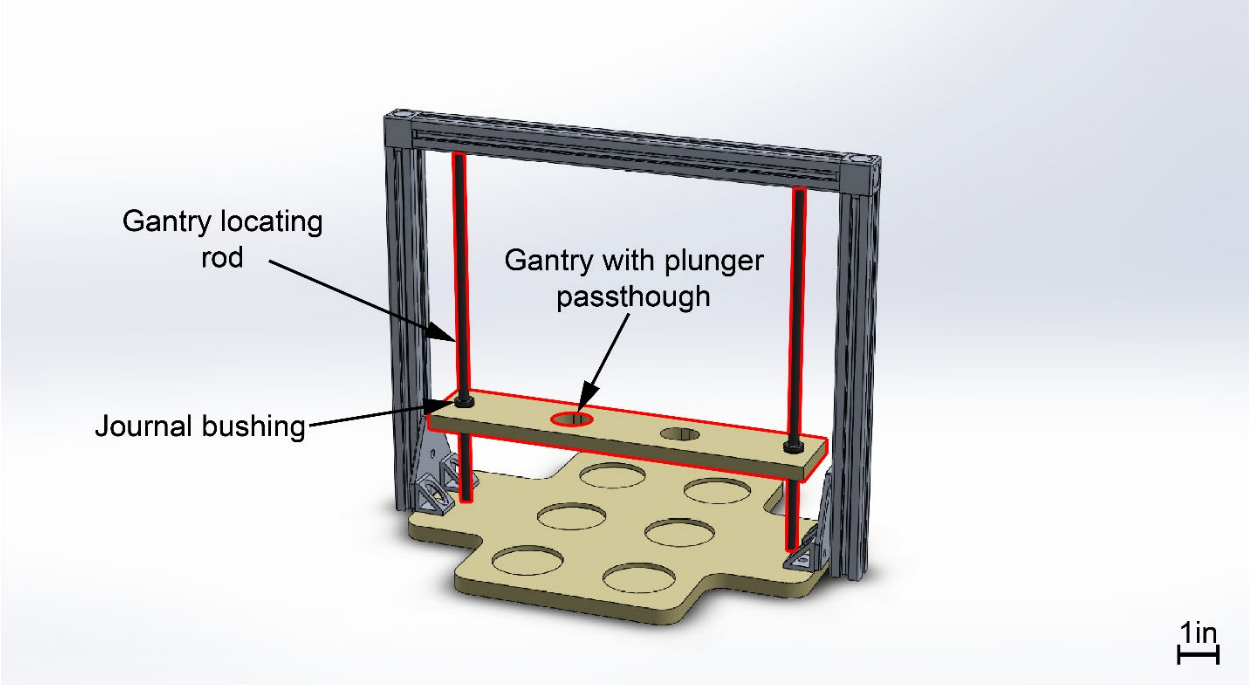


*Figure S3: The polysulfone gantry is suspended and held in alignment with journal bushings, allowing unrestricted vertical motion. Plunger passthroughs are machined inside the gantry to allow the clearance of two center plungers.*

The locating rods were initially 316 stainless steel, but after months of operation in high temperature and relative humidity, the initial stainless rods were replaced with anodized 6061 aluminum to address surface corrosion. Low friction PTFE journal bushings were press-fit into the gantry and used to provide interface between the gantry and locating rods. To connect the polysulfone gantry to the superstructure, a central motor mount was connected (Figure S4).

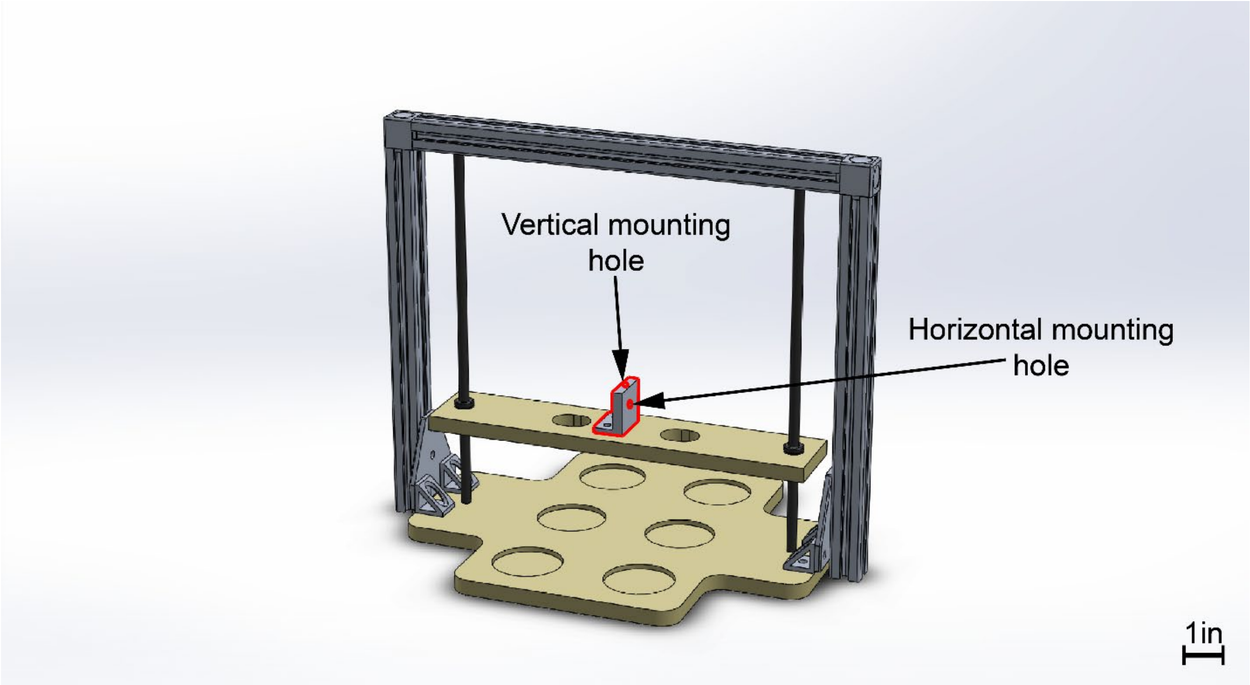


*Figure S4: Aluminum center motor mount features points for both the horizontal and vertical motor connections. This motor mount serves to connect the gantry to the superstructure.*

Since the entire gantry subassembly will be the focus of PID tuning later, the components which attach to it should be kept at the lowest weight possible to aid in ease of tuning. The central motor mount was machined from aluminum and kept relatively thin to keep weight low. A large aluminum plunger plate is then rested on top of the gantry (Figure S5).

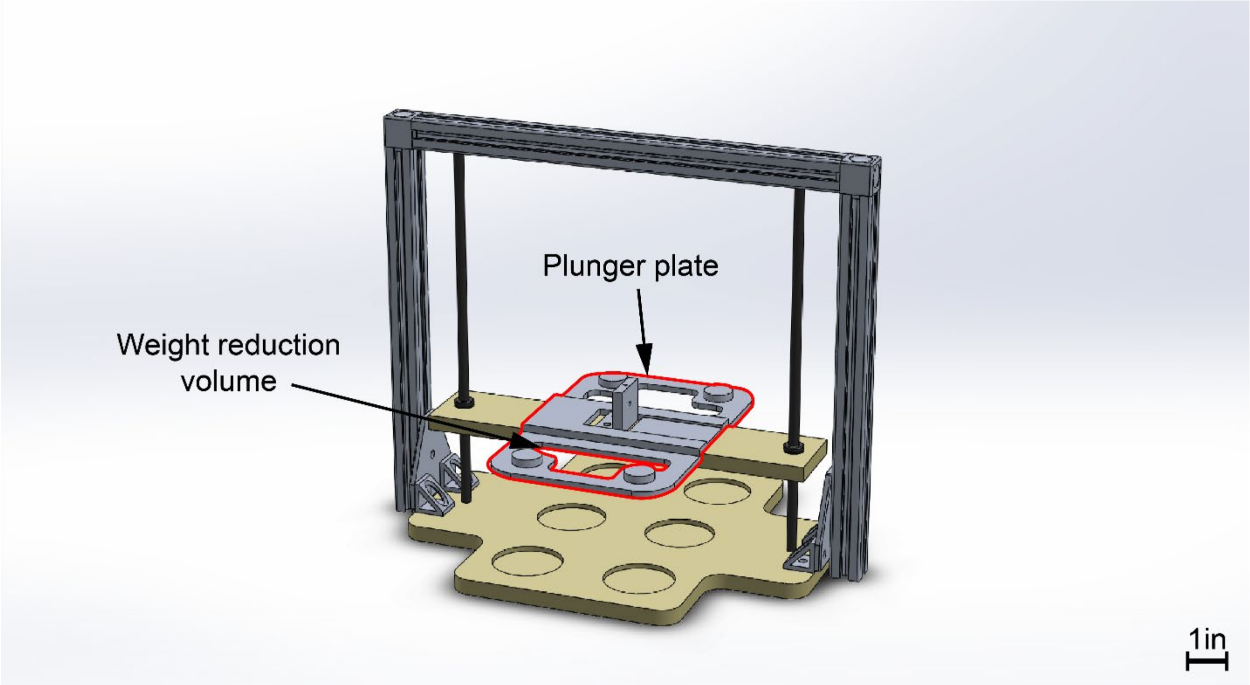


*Figure S5: Aluminum plunger plate slides freely on the upper surface of the gantry. Six reinforced plate sections are threaded to receive plunger screws.*

This aluminum plate serves the primary role of suspending polysulfone plungers above individual wells within the base plate. Material was removed from the plunger plate to allow retention of strength while shedding as much weight as possible. The plunger plate uses a recessed channel to limit the degrees of motion relative to the gantry, allowing sliding motion on one axis only. Plungers were then attached to the underside of the plate (Figure S6).

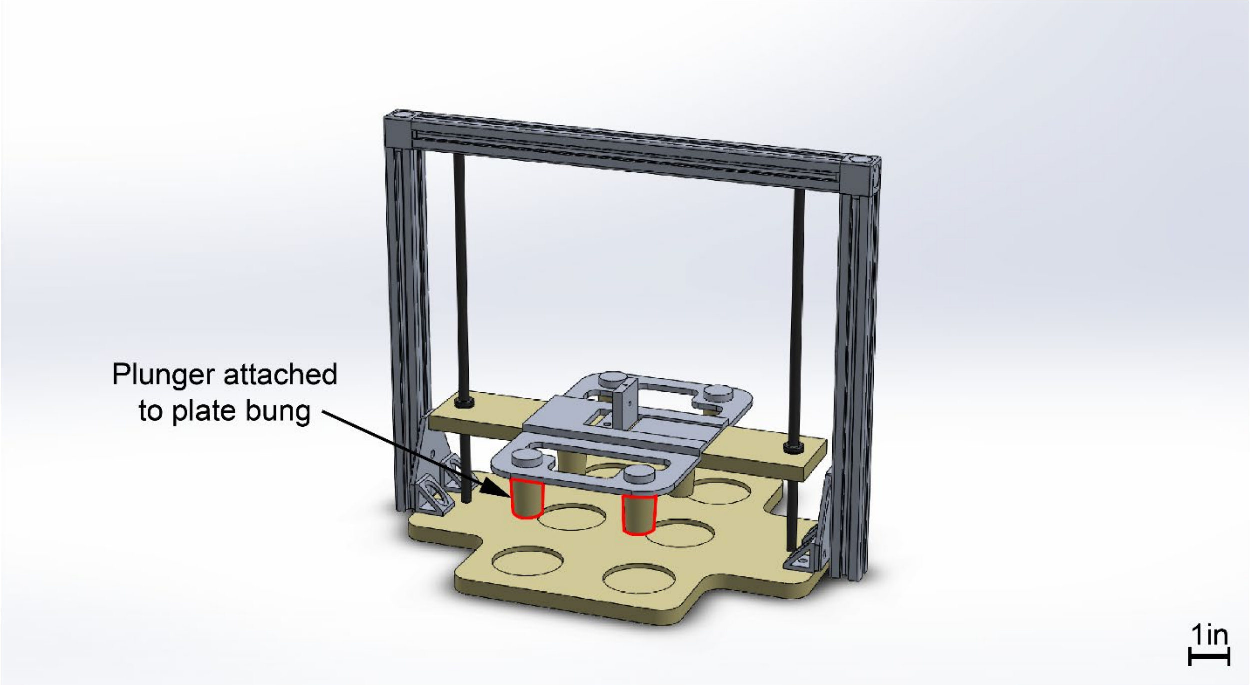

*FigureS6: Polysulfone plungers hang from the plunger plate, with each plunger aligning with a single base plate recess.*

The polysulfone plungers serve as the point of contact with samples during loading experiments, and as such need to be simultaneously faced for uniform length after individual plugs were turned on a lathe. To attach the actuation motors to the superstructure/gantry, a horizontal and vertical motor mount were machined and mounted. The horizontal motor mount was fastened to the plunger plate (Figure S7).

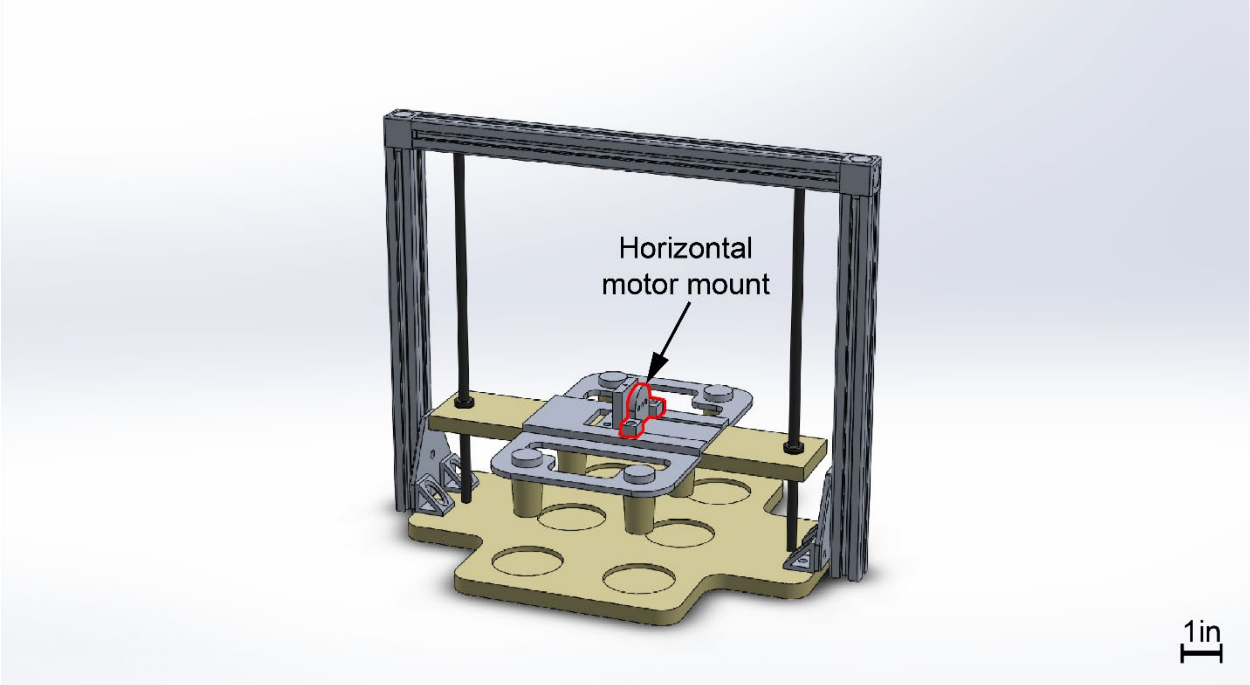


*Figure S7: Horizontal motor mount creates an attachment point between the gantry and plungers.*

Next, the horizontal motor could be secured to the horizontal motor mount, and its output shaft secured to the center motor mount. This effectively connects the gantry to the plunger plate (Figure S8).

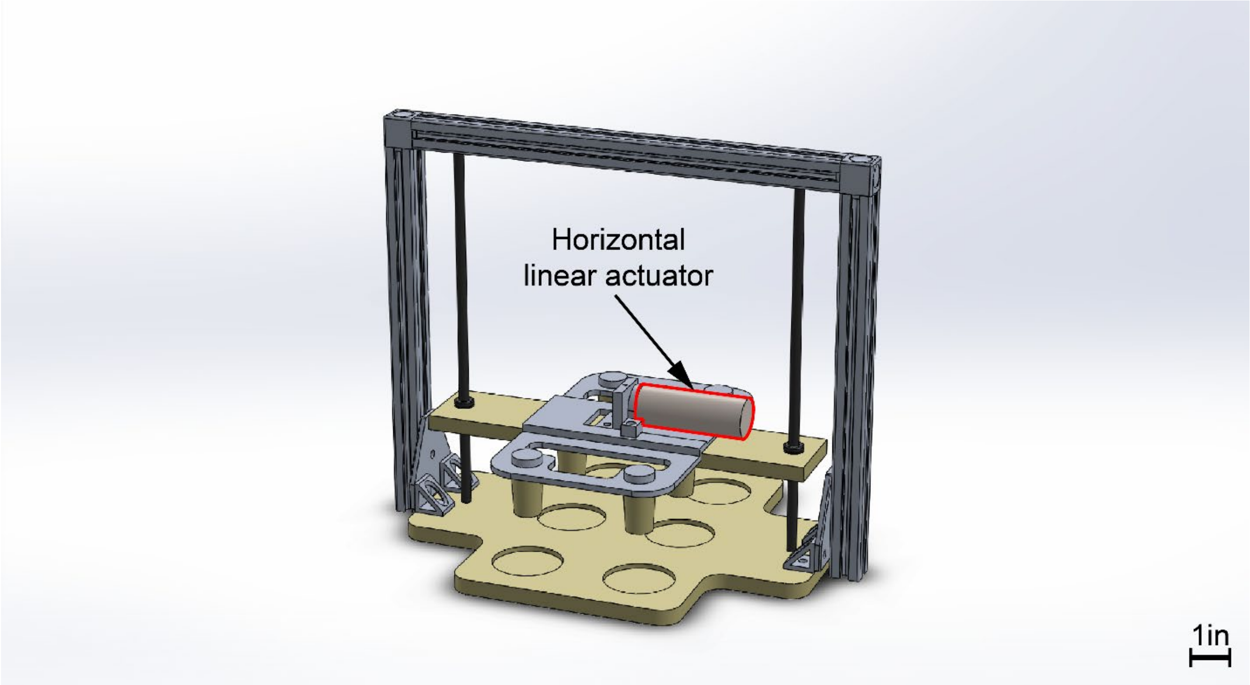


*Figure S8: Horizontal linear actuator forming the connection between gantry and plunger plate.*

To connect the gantry to the superstructure, the vertical motor had to be hung above the center motor mount (Figure S9).

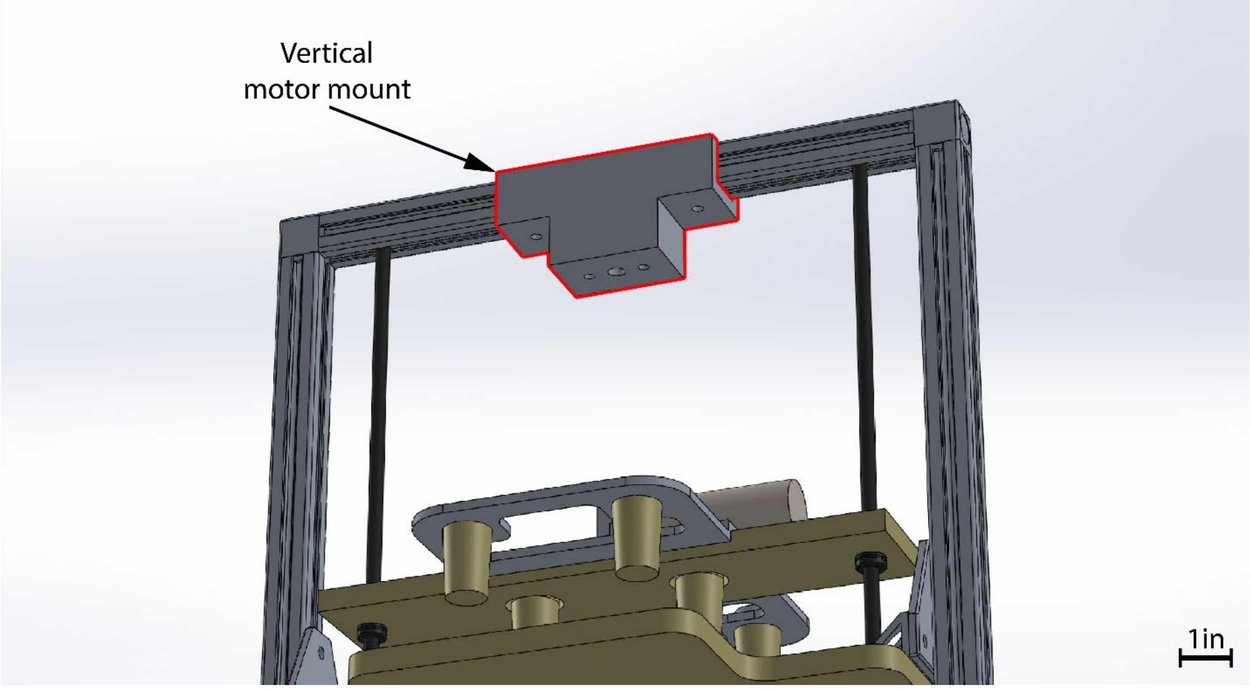


*Figure S9: Vertical motor mount located the vertical motor above the center motor mount.*

The vertical linear actuator could then be suspended between the superstructure and the gantry (Figure S10).

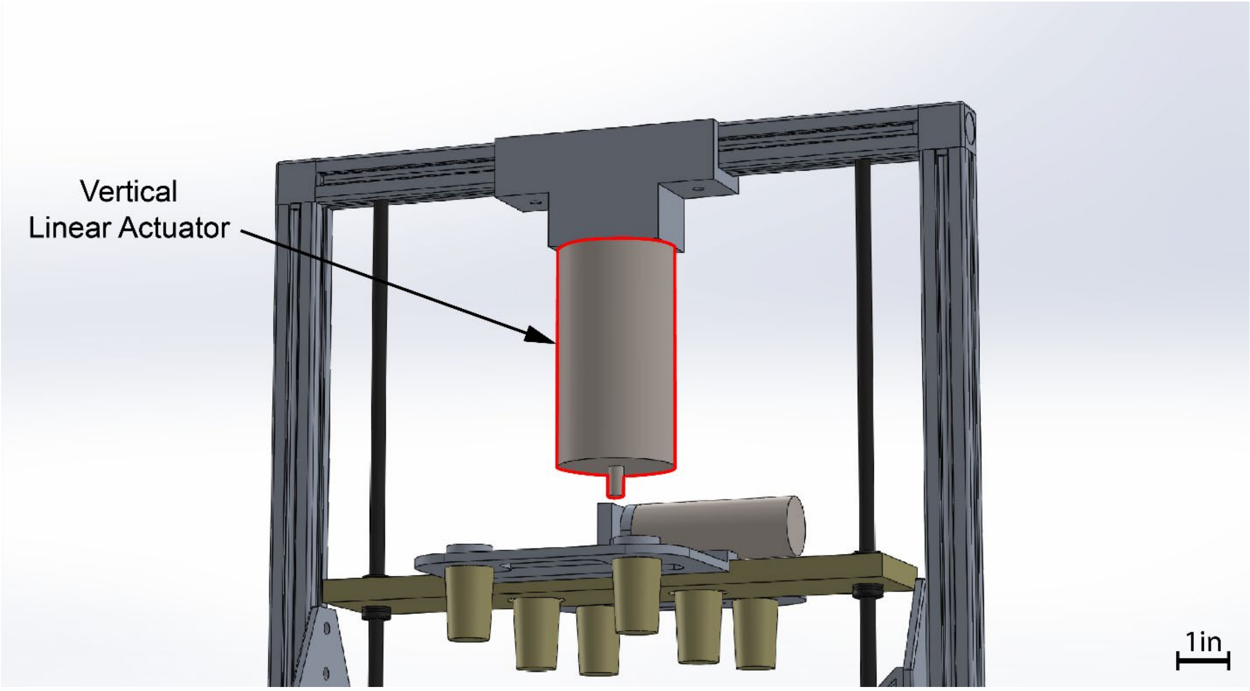


*Figure S10: Vertical motor positioned above the gantry and aligned with the center motor mount.*

Finally, the vertical motor output shaft is secured to the center motor mount, creating the connection between superstructure, gantry, and plunger plate (Figure S11). To facilitate ease of assembly, the connection between the center mount and vertical motor shaft is made using a brass rod which had both a positive and negative thread pitch, allowing the connector to be threaded into both the shaft and center motor mount simultaneously.

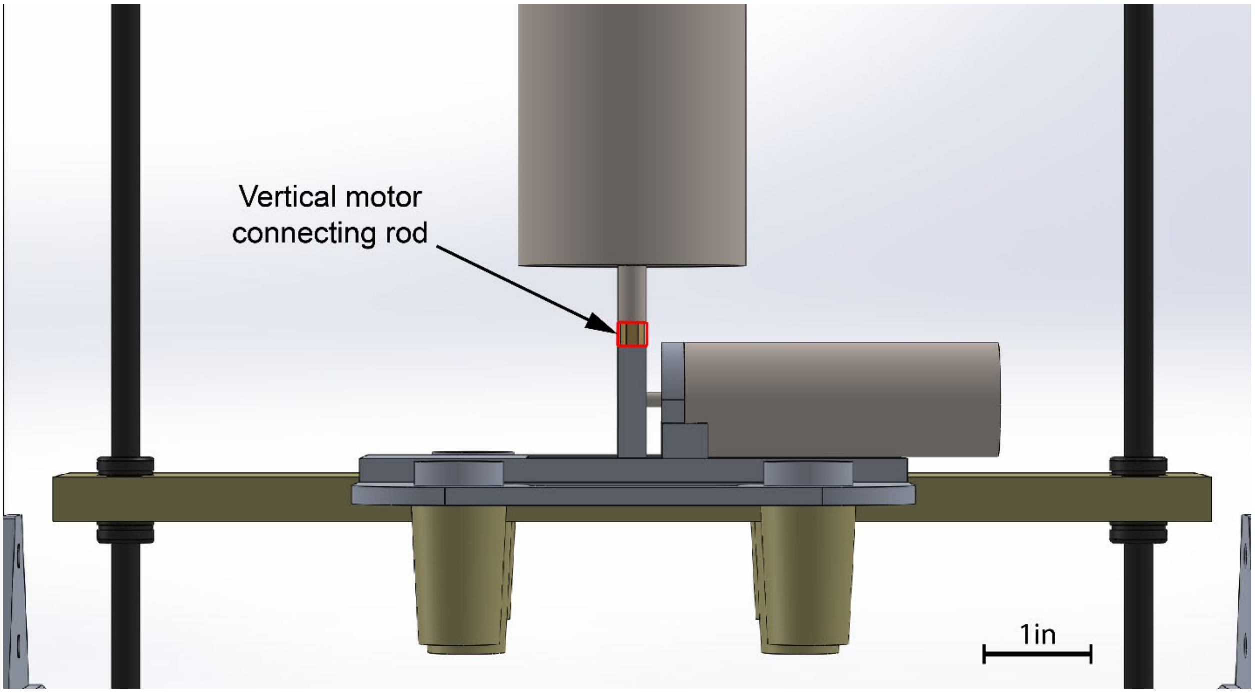


*Figure S11: Motor connections made at the center motor mount, including the brass dual-thread center connection.*

Once the motor shafts have been connected to the center motor mount, the machine assembly is completed (Figure S12).

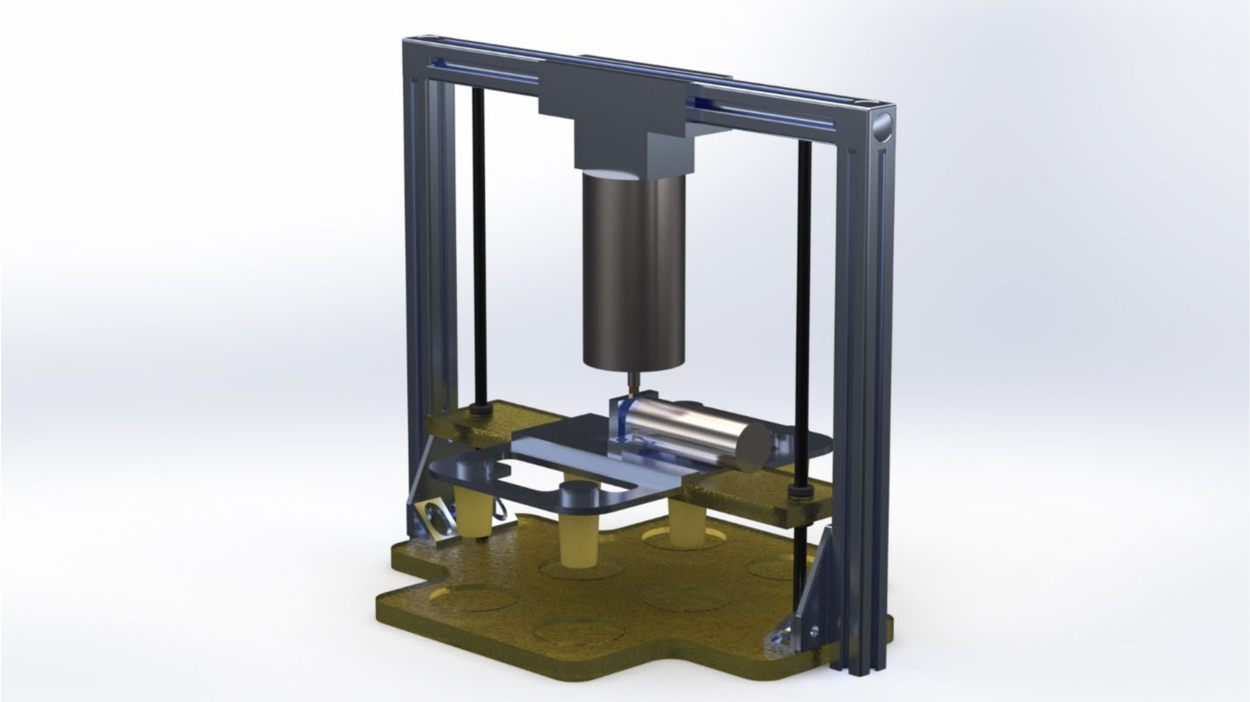

*Figure S12: Render of fully assembled machine, showing differences in component material and assembly orientation.*

When completed, the entire assembled machine weighs 8.2 pounds. Overall machining practices followed the expected design without notable exception. The final render of the SolidWorks model in Figure S12 closely resembles the finalized design in layout, function, size, and weight (Figure S13).

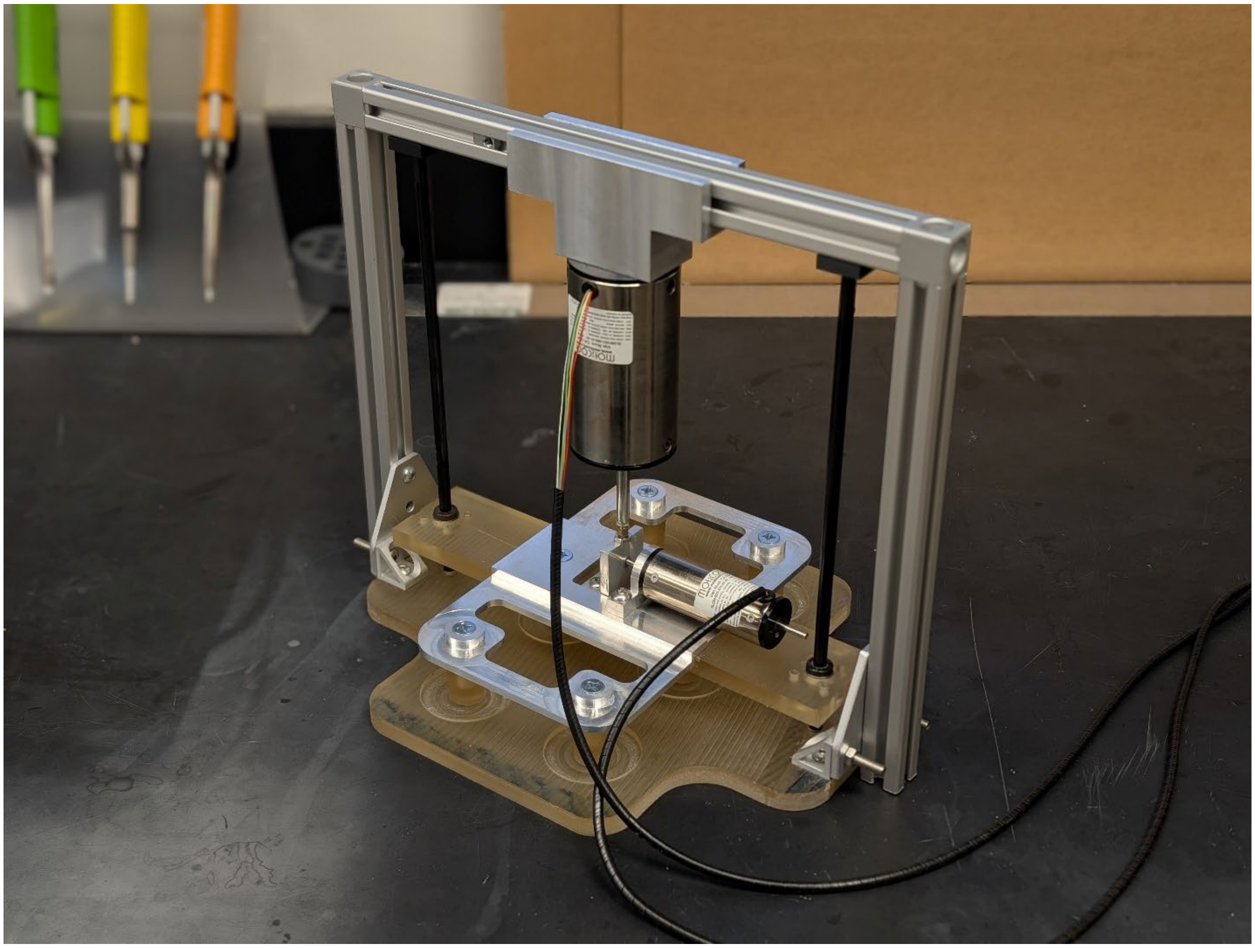

*Figure S13: Machined and assembled bioreactor with Moticont linear actuators closely resembles expected design.*

To actuate the gantry and plunger plate, linear actuators were selected which suited both the necessary actuation force and encoder resolution (Moticont, Van Nuys, CA). Drawings representing each of the manufactured bioreactor components are available below.

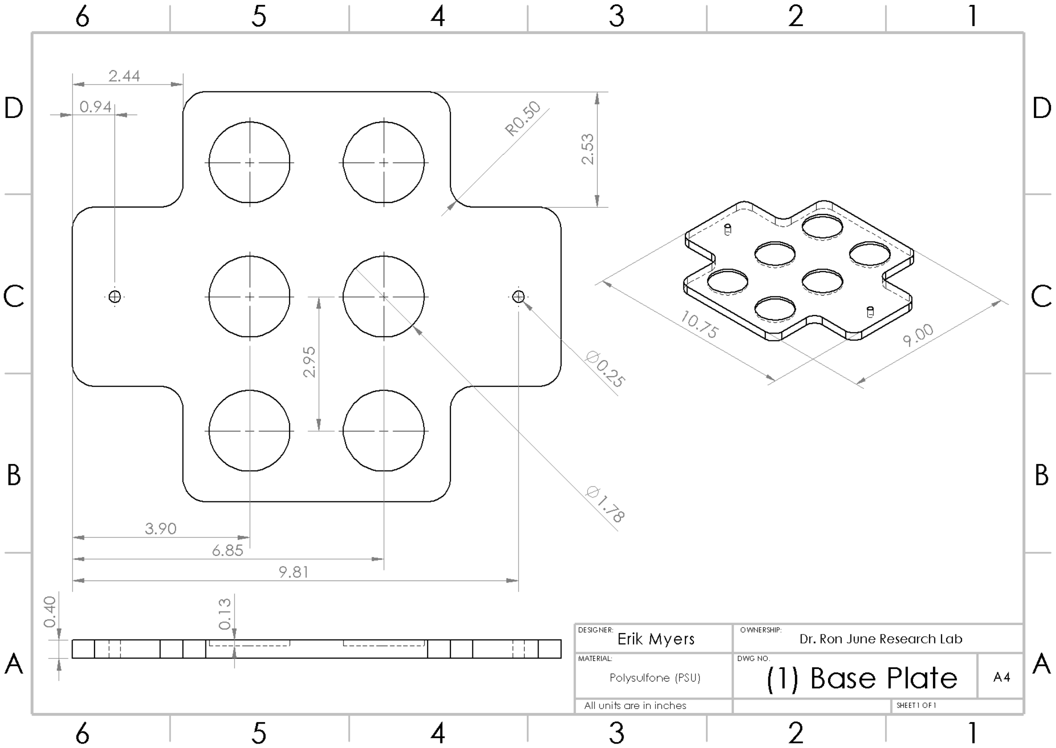
2.44
0.94
R0.50
2.53
2.95
Ø0.25
Ø1.78
3.90
6.85
9.81
0.40
0.13
10.75
9.00
DESIGNER: Erik Myers
OWNERSHIP: Dr. Ron June Research Lab
MATERIAL: Polysulfone (PSU)
DWG NO. (1) Base Plate
A4
All units are in inches
SHEET 1 OF 1

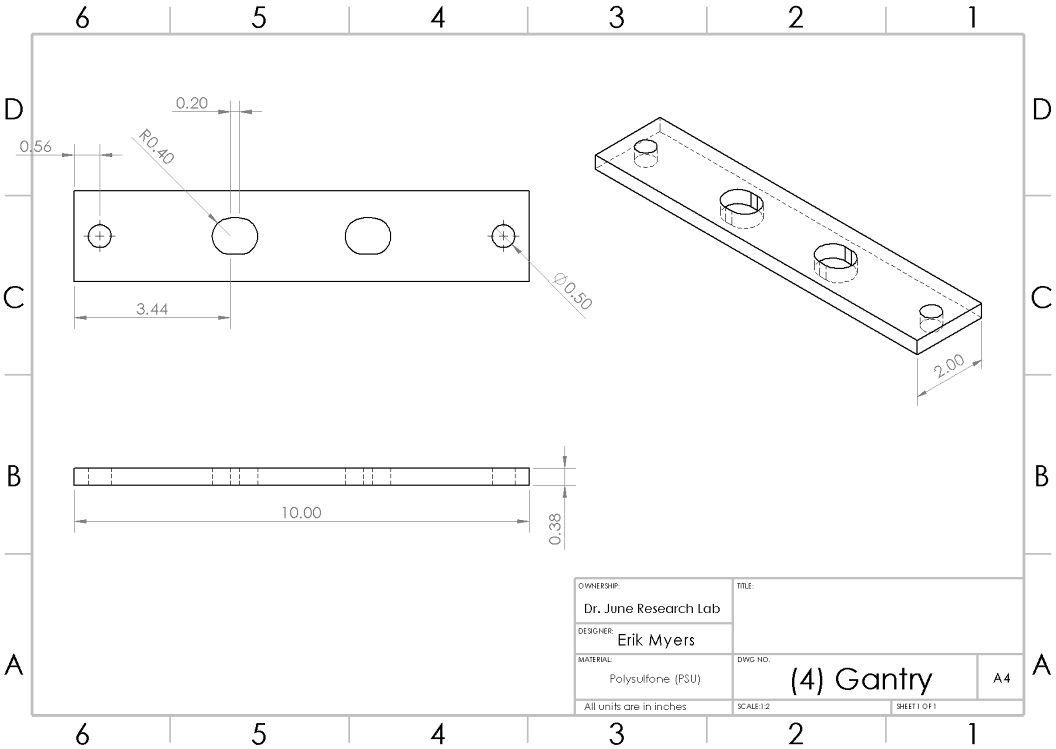
0.20
0.56
R0.40
Ø0.50
3.44
2.00
10.00
0.38
OWNERSHIP: Dr. June Research Lab
TITLE:
DESIGNER: Erik Myers
MATERIAL: Polysulfone (PSU)
DWG NO. (4) Gantry
A4
All units are in inches
SCALE:1:2
SHEET 1 OF 1

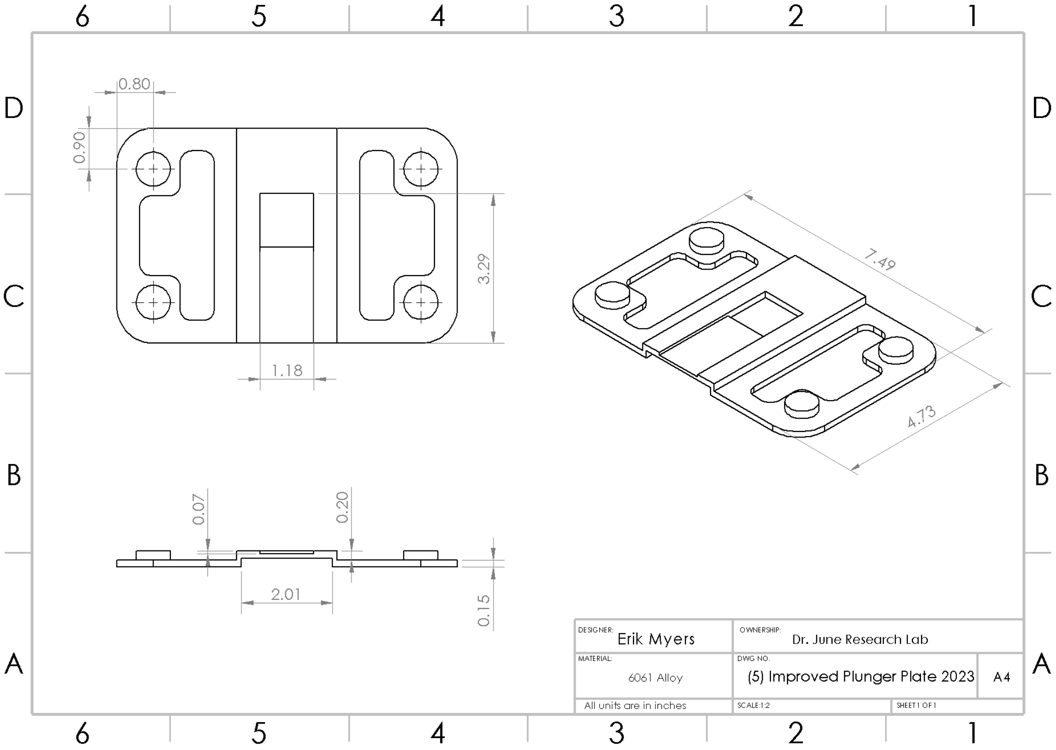
0.80
0.90
3.29
1.18
7.49
4.73
0.07
0.20
2.01
0.15
DESIGNER:
Erik Myers
OWNERSHIP:
Dr. June Research Lab
MATERIAL:
6061 Alloy
DWG NO.
(5) Improved Plunger Plate 2023
A4
All units are in inches
SCALE:1:2
SHEET 1 OF 1

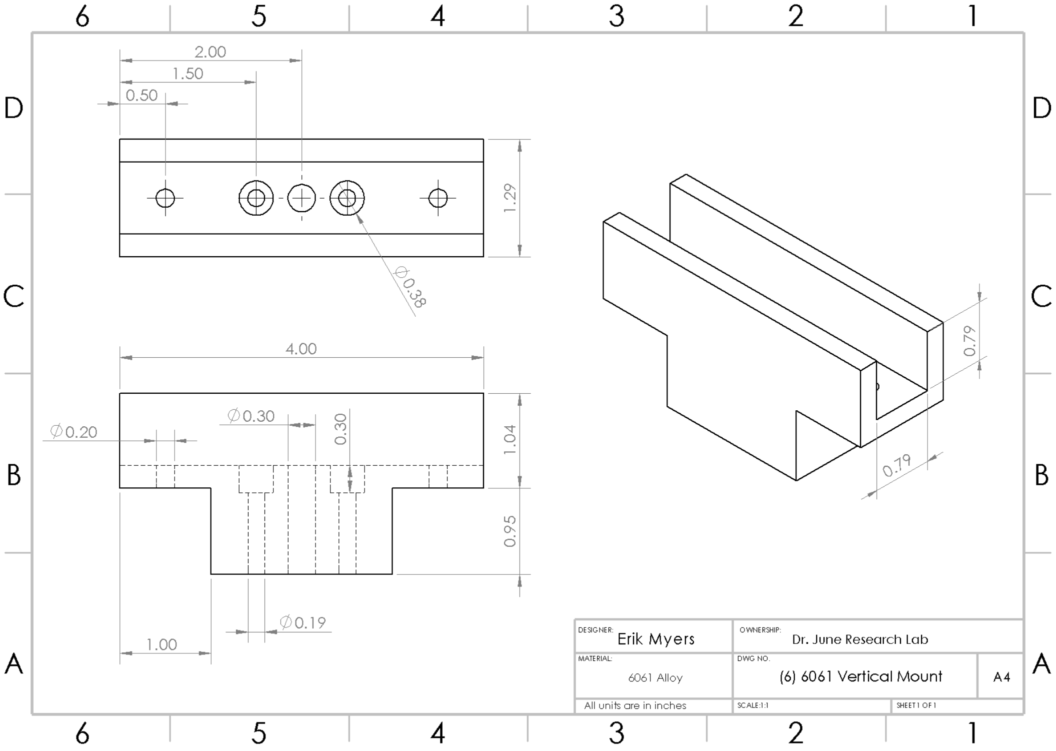
2.00
1.50
0.50
1.29
∅0.38
4.00
∅0.30
0.30
∅0.20
1.04
0.95
∅0.19
1.00
0.79
0.79
DESIGNER:
Erik Myers
OWNERSHIP:
Dr. June Research Lab
MATERIAL:
6061 Alloy
DWG NO.
(6) 6061 Vertical Mount
A4
All units are in inches
SCALE:1:1
SHEET 1 OF 1

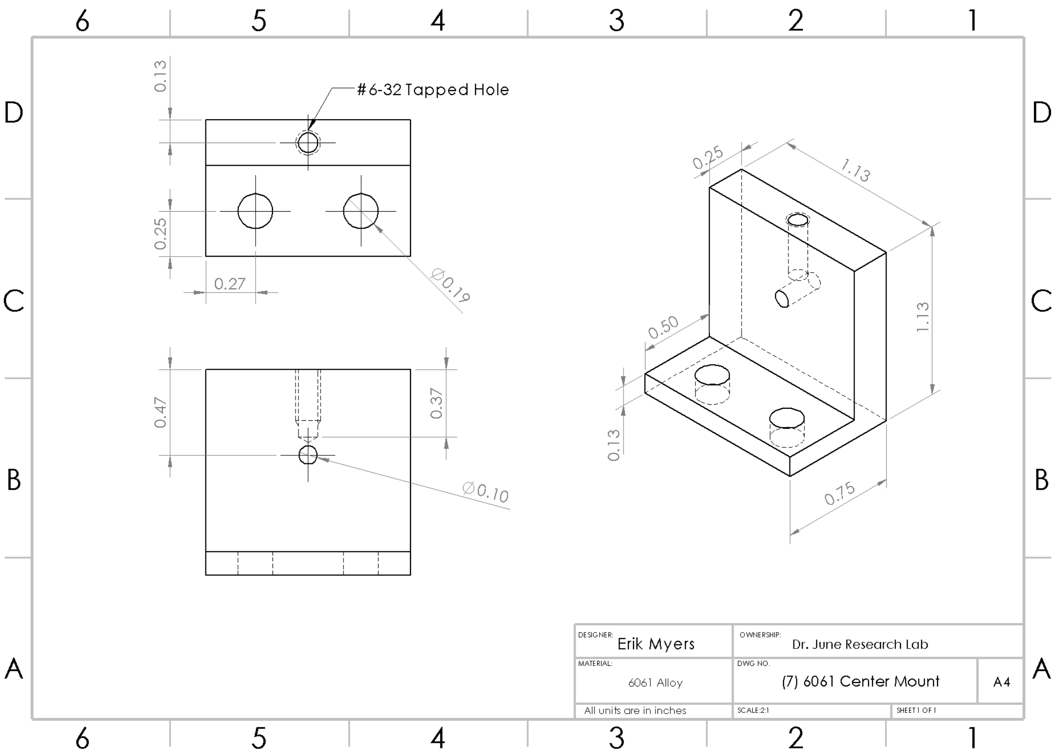
#6-32 Tapped Hole
0.13
0.25
0.27
∅0.19
0.47
0.37
∅0.10
0.25
1.13
1.13
0.50
0.13
0.75
DESIGNER: Erik Myers
OWNERSHIP: Dr. June Research Lab
MATERIAL: 6061 Alloy
DWG NO. (7) 6061 Center Mount
A4
All units are in inches
SCALE:2:1
SHEET 1 OF 1

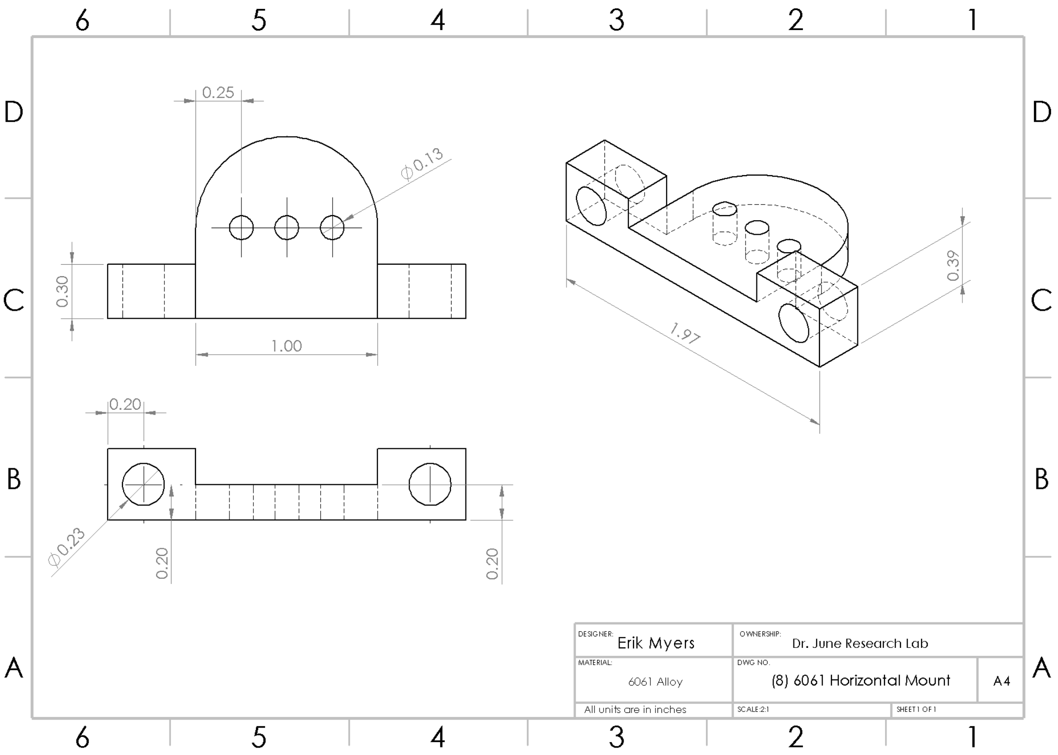
0.25
∅0.13
0.30
1.00
0.39
1.97
0.20
∅0.23
0.20
0.20
DESIGNER: Erik Myers
OWNERSHIP: Dr. June Research Lab
MATERIAL: 6061 Alloy
DWG NO. (8) 6061 Horizontal Mount
A4
All units are in inches
SCALE:2:1
SHEET 1 OF 1

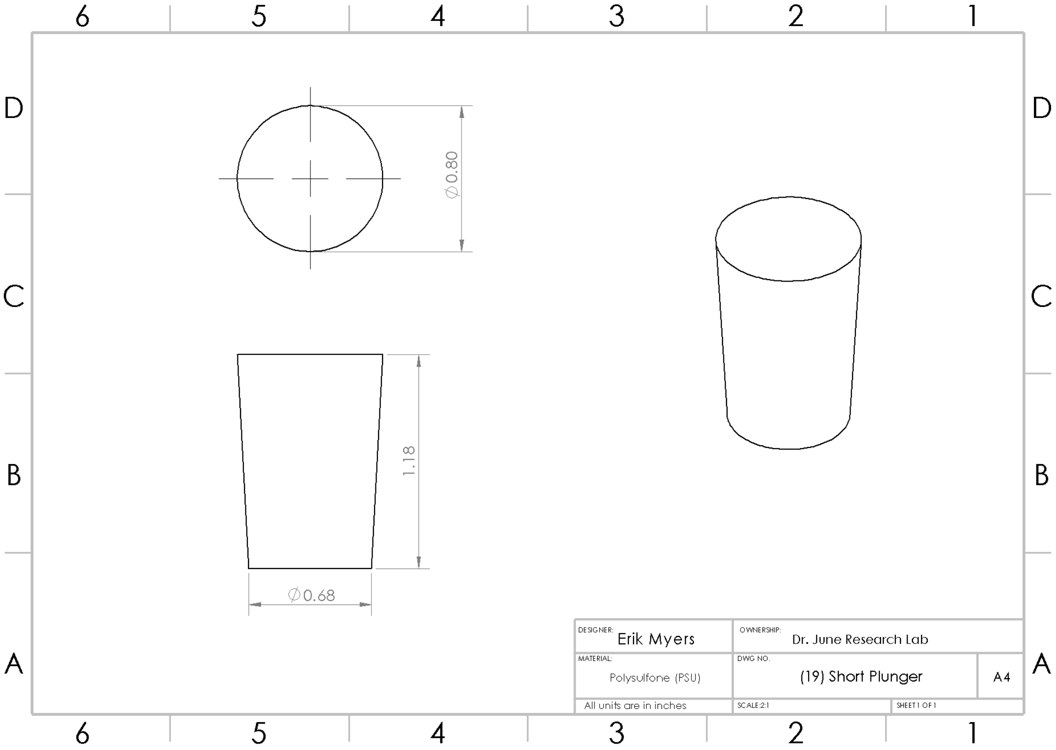

Ø0.80
1.18
Ø0.68
DESIGNER: Erik Myers
OWNERSHIP: Dr. June Research Lab
MATERIAL: Polysulfone (PSU)
DWG NO. (19) Short Plunger
A4
All units are in inches
SCALE 2:1
SHEET 1 OF 1
