## Supplemental File 2 for "Targeted analysis of chondrocyte central metabolites in response to cyclical compression and shear deformations"

### Supplemental File 2: Extraction Optimization and Outlier Analysis

During the process of metabolite extraction, it was discovered that the previously used protocol extraction buffer (70:30:1% Methanol:Acetone:HCl) created a film in the supernatant which could clog the electro-spray ionization nozzle within the mass spectrometry machine. After the extraction of the first four donors with this method, the buffer was switched to the final buffer (70:30:1% Methanol:Acetone:Formic acid). Unfortunately, Donor #1 was unable to be cleaned substantially with the use of acetonitrile dilutions and column filtering and were subsequently removed from the data set. Donors #3 and #4 were able to be rescued using the same technique of HPLC Water:Acetonitrile dilution and column filtering with centrifugal spinning, but the results of their mass spectrometry analysis differ from the remaining 12 donors in particular metabolite concentrations.

Since three of the remaining 14 samples were extracted using a different extraction buffer (Donors 2,3, and 4), a preliminary analysis was performed using MetaboAnalyst to discover if there were concerning differences in the reported results from the mass spectrometry analysis. A Principal Component Analysis (PCA) was performed that showed immediate obvious separations between groups (Figure S2.1).

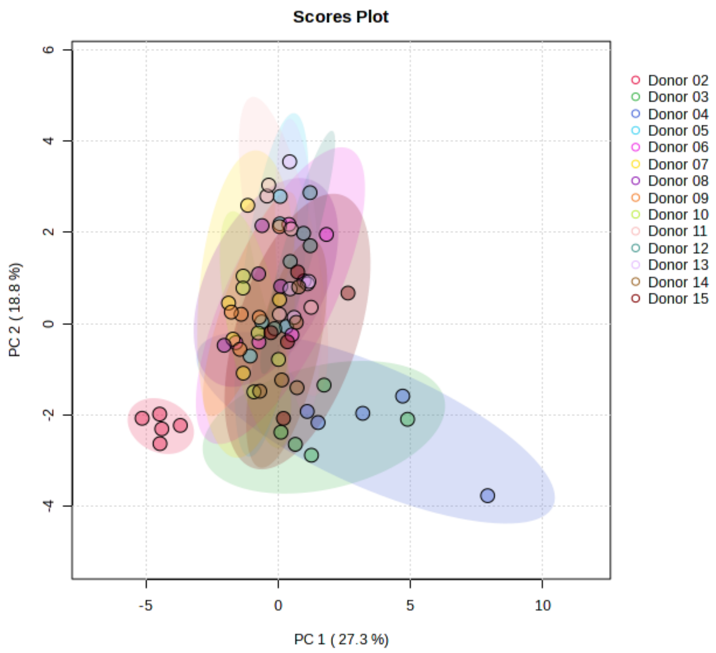


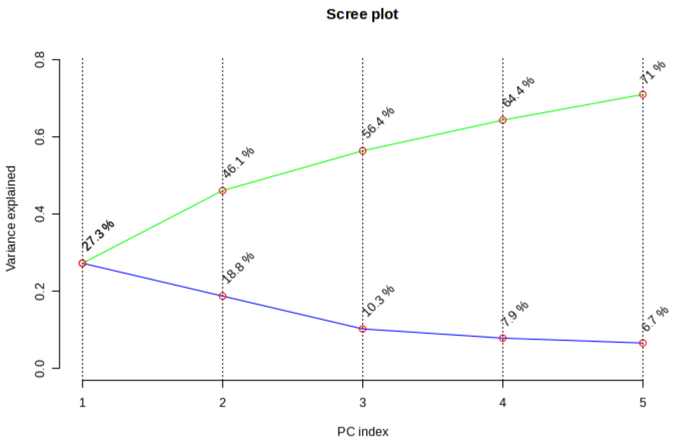


*Figure S2.1: (Top) PCA analysis on a per-donor basis revealed differences between Donor 2 and the remaining donor conglomerate. Donors 3 and 4 also showed some separation but were included in the study due to significant overlap in groupings. (Bottom) Scree plot showing contribution of principal components, with 46.1% of variance being explained in the first two components.*

Donor 2 showed clear differences when compared to other groups in the first two principal components, which explains 46.1% of the variance in the donor data sets. Because of this, datum from Donor 2 was removed from the dataset. This is supported by clustering analysis where the dendrogram of the donor groupings also finds that Donor 2 was clearly separated from the remaining donors (Figure S2.2).

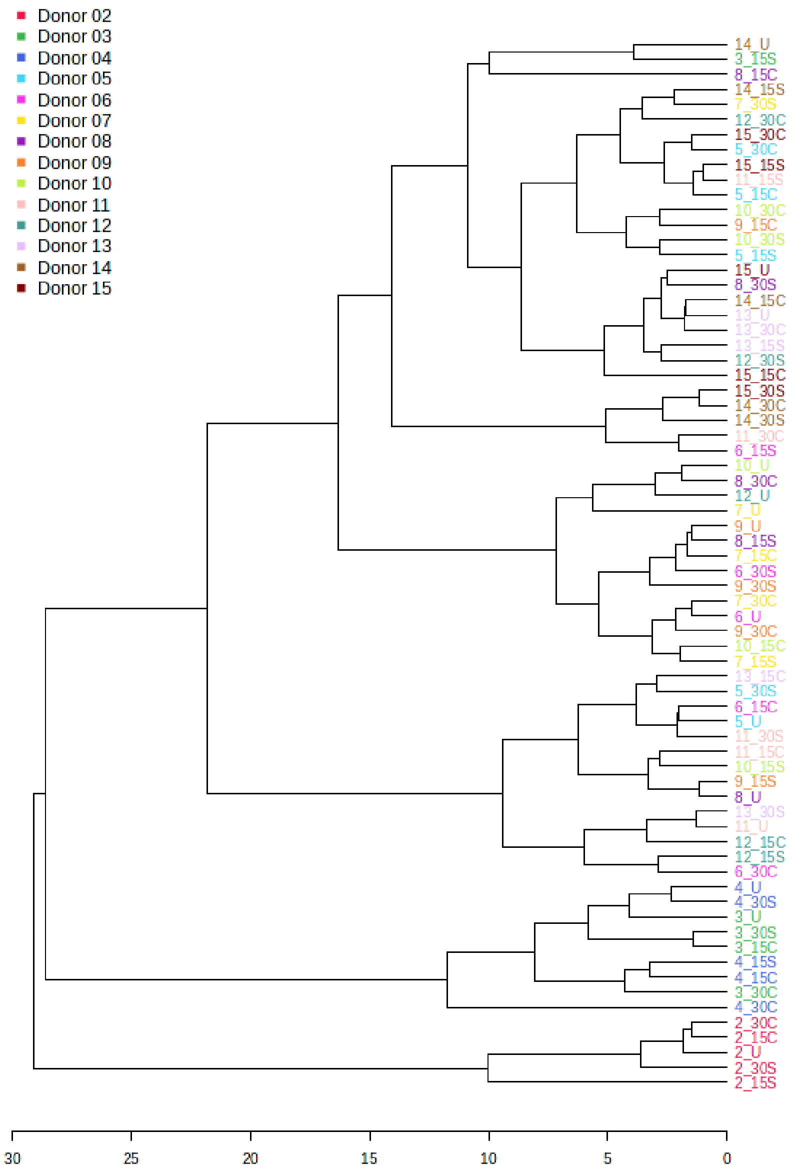


*Figure S2.1: Donor 2 not only grouped exclusively with itself, but also showed greater magnitude of separation than any other grouping in the dendrogram.*

Once Donor 2 was removed from the dataset, this analysis was re-run to see the apparent differences in donors 2 and 3 from the remaining 13 donors of the dataset (Figure S2.3).

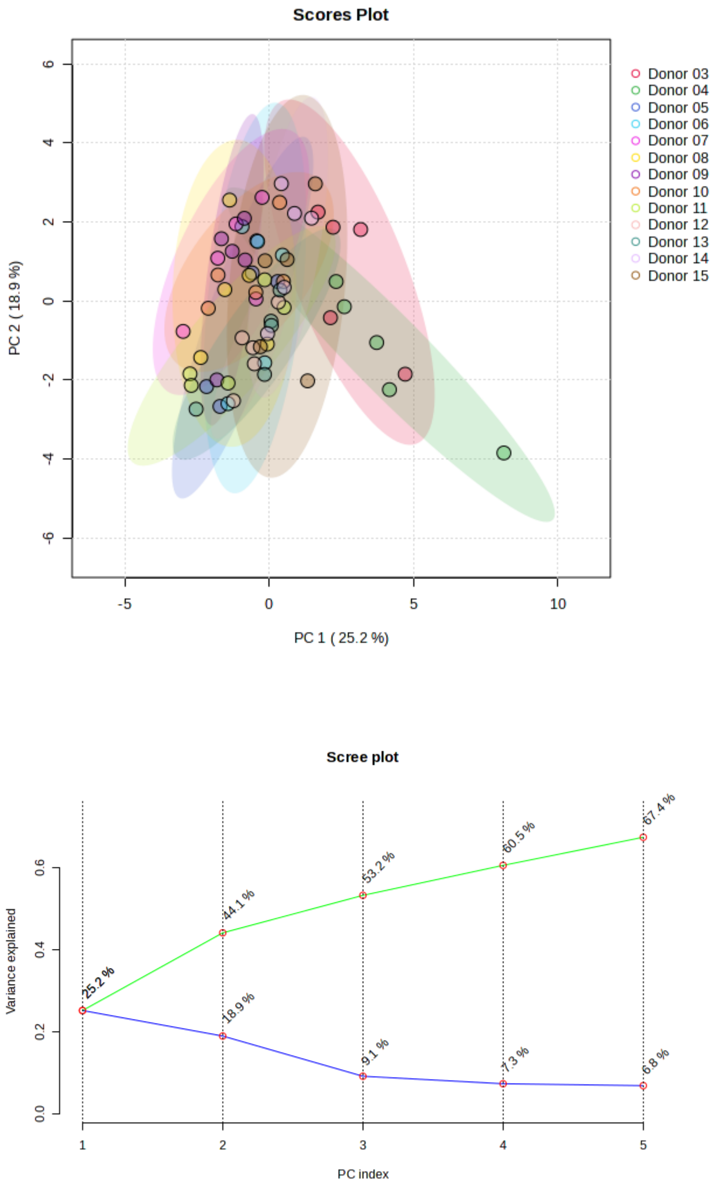


*Figure S2.2: (Top) PCA of trimmed dataset shows separation of donors 3 and 4, but without a clear independent grouping to warrant removal from the dataset. (Bottom) Scree plot still shows a large contribution from the first 5 principal components, with 44.1% of the variance being explained with the first two components.*

Donors 3 and 4 showed greater separation from the other responses once donor 2 was removed, but the resulting dendrogram suggested the differences were not as pronounced (Figure S2.4).

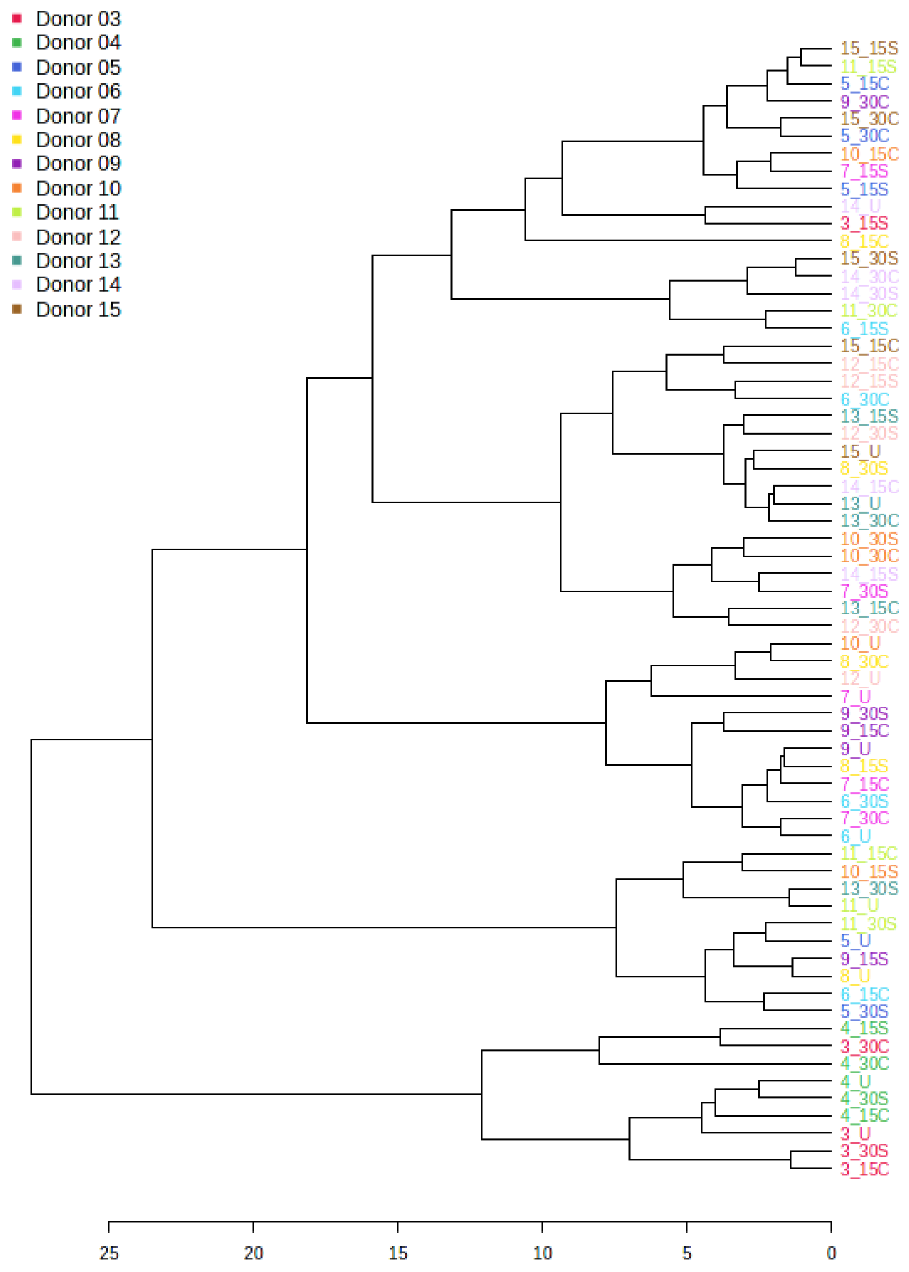


*FigureS2.3: Dendrogram of the trimmed dataset shows grouping of donors 3 and 4. However, the Euclidian distance is relatively small in comparison to the separation seen by donor 2, suggesting that donors 3 and 4 may not be different enough to remove from the dataset.*

Given the results of the initial donor PCA and dendrograms, the decision was made to exclude donor 2 from the dataset moving forward. This is the trimmed dataset with N=13 bovine and human donors.

Since 1/3 of the initial donor set was comprised of bovine cells (Donors 11-15), it was necessary to assess the effect of species on data grouping. If there was enough evidence to suggest that human and bovine donors reacted differently to mechanical stimulation, further statistical analyses would require species separation. Again, a PCA was performed to explore differences in metabolite profiles in a two-dimensional projection based on species (Figure S2.5).

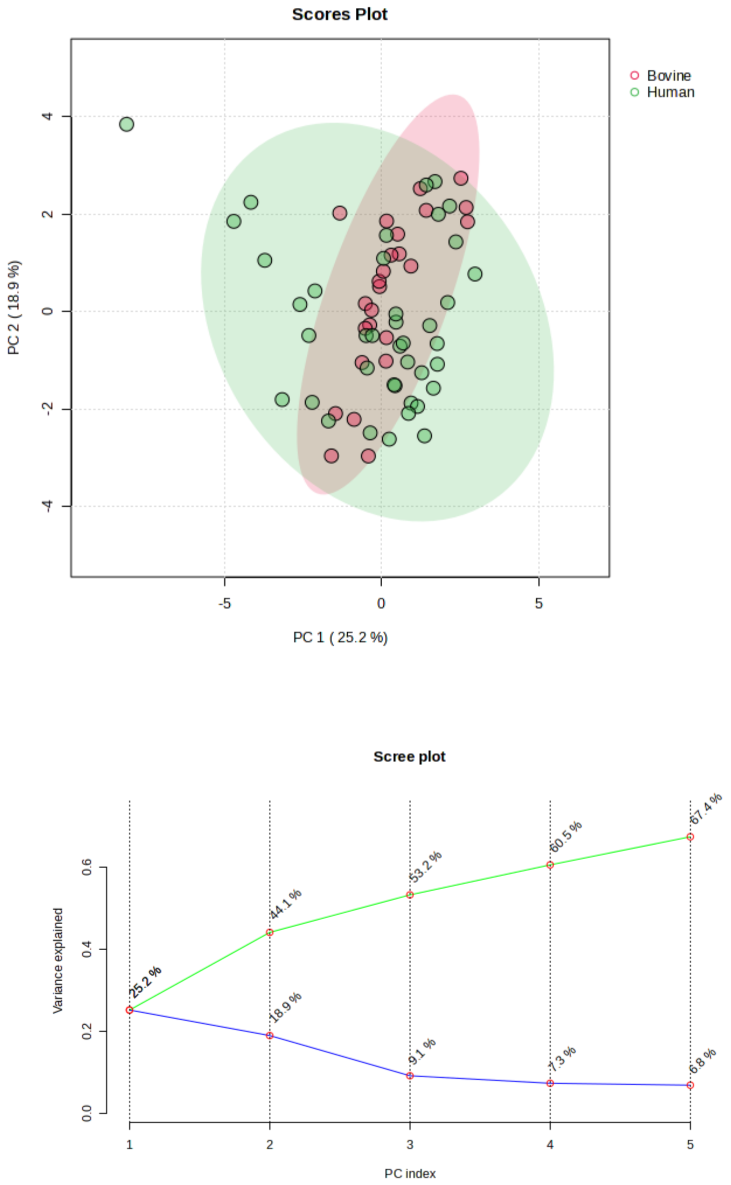


*FigureS2.4: (Top) PCA analysis of species-based variation revealed considerable overlap between human and bovine donors, with 44.1% of the total variance explained in the first two components. (Bottom) Scree plot shows percentage of variation explained by five principal components.*

While Figure S2.5 did demonstrate different variation in the first PC the overlap in the variance of human samples when compared to bovine donors, the overlap of groups suggests that there is not a difference between human and bovine donors. The larger variance of the human samples in the first PC is most likely because human donors are different in many aspects including age, race, sex, weight, and OA progression at the time of joint donation. Bovine donors, for the most part, are very uniform in age, sex, weight, and health at the time of donation. Regardless, the results of the PCA suggest that human and bovine donors can be treated as similar in future statistical analyses.

Before the metabolite-specific statistical analyses could be performed, final analyses were performed and differences based on the donor gender in the metabolic response profiles were explored to determine if there were effects on metabolic response based on donor gender (Figure S2.6).

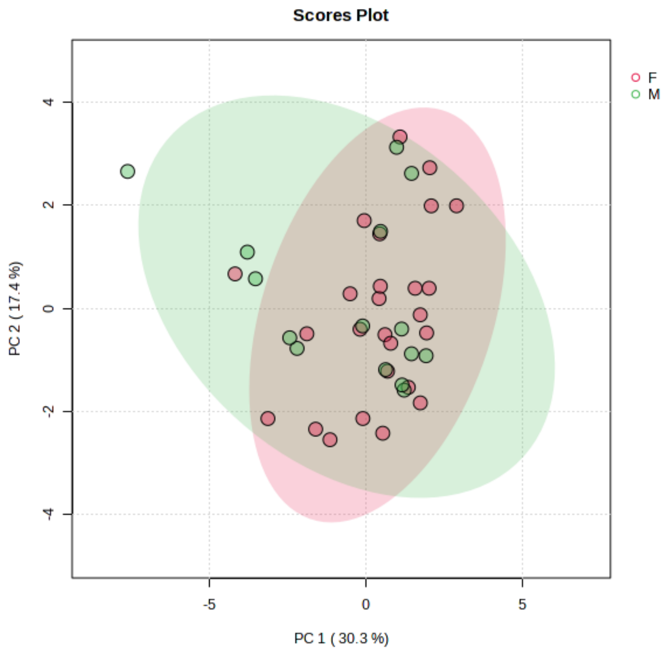

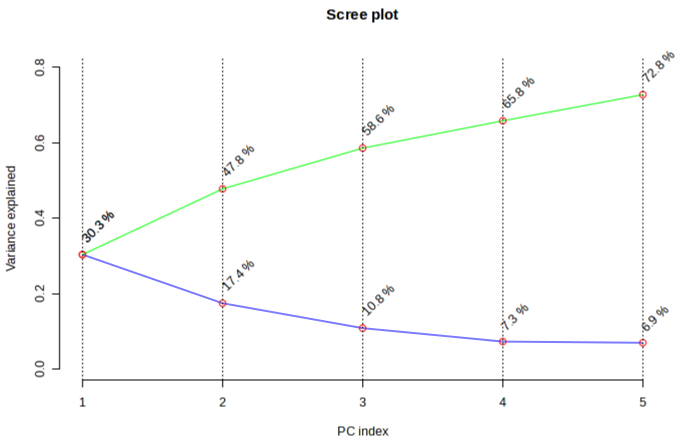


*Figure S2.5: (Top) Gender-based PCA showed similarities in variation between male and female donors. (Bottom) Scree plot shows even greater contribution of first two principal components, explaining 47.8% of total variance.*

It should be noted that the gender-based comparison was performed only with the remaining nine human donors, since the sex of bovine donors is unknown. A median heatmap was also used to check for trends in differences between male and female donors for metabolite concentrations.

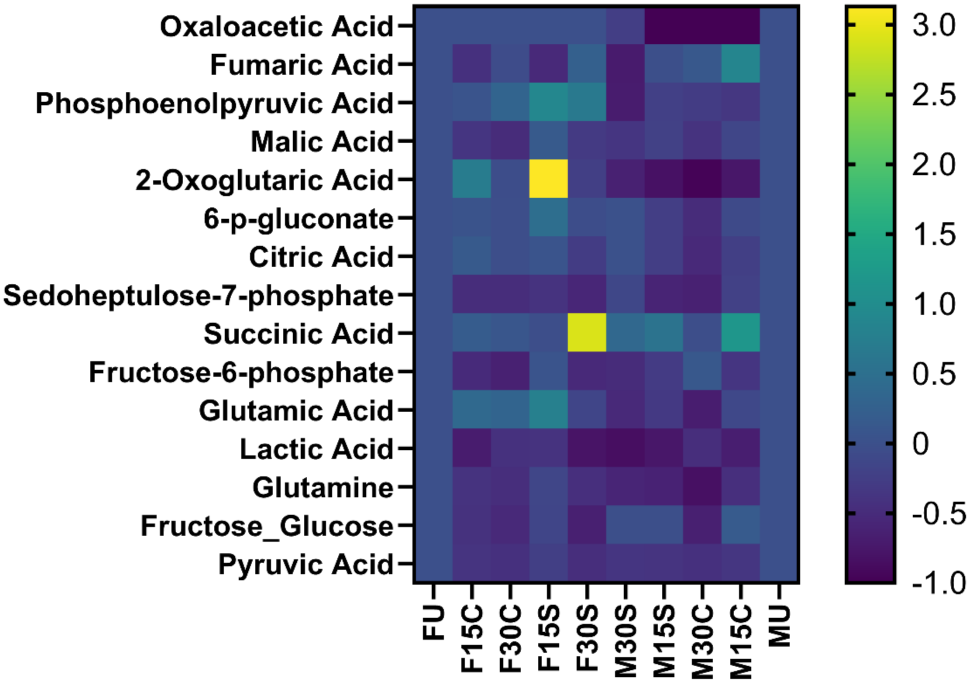


*Figure S2.7: Median heatmap of group metabolite intensities separated by sex reveals no concerning differences between male and female. Group median intensities are normalized versus the unloaded control intensity. Oxaloacetate showed decreased median values for all male donors, but this trend is due to the very poor signal quality of oxaloacetate.*

Based on the evidence from Figures S2.5, S2.6, and S2.7, it was decided to perform individual metabolite statistics with all male human, female human, and bovine samples together.
