## Supplemental File 3 for "Targeted analysis of chondrocyte central metabolites in response to cyclical compression and shear deformations"

### Supplemental File 3: Mass Spectrometry Calibration Data

#### Quantitative Analysis Calibration Report

| | | | |
|---|---|---|---|
| **Batch Data Path** | J:\6538\shared\20241016_01_Eric_TCAcells_BEHC18-100\QuantResults\20241016_EMPB.batch.bin | | |
| **Analysis Time Stamp** | 10/17/2024 2:51:43 PM | **Analyst Name** | MSU\w68p731 |
| **Report Generation Time** | 10/17/2024 3:01:12 PM | **Reporter Name** | MSU\w68p731 |
| **Calibration Last Update** | 10/17/2024 2:44:43 PM | **Batch State** | Processed |
| **Analyze Quant Version** | 12.0 | **Report Quant Version** | 12.0 |

**Oxaloacetic acid** **Relative Standard Error** 19.5

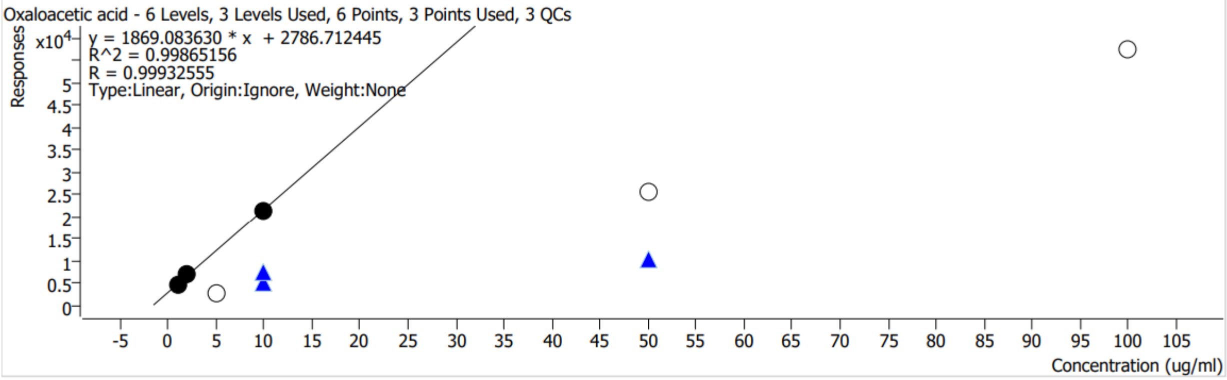


| Calibration STD Path | Type | Level | Enable | Conc. | Response | RF | Level RSD |
|---|---|---|---|---|---|---|---|
| J:\6538\shared\20241016_01_Eric_TCAcells_BEHC18-100\20241016_EMPB__05_1_Std.d | Calibration | 1 | x | 1.0000 | 4339 | 4338.5718 | |
| J:\6538\shared\20241016_01_Eric_TCAcells_BEHC18-100\20241016_EMPB__05_2_Std.d | Calibration | 2 | x | 2.0000 | 6882 | 3440.8785 | |
| J:\6538\shared\20241016_01_Eric_TCAcells_BEHC18-100\20241016_EMPB__05_5_Std.d | Calibration | 5 | | 5.0000 | 2830 | 565.9365 | |
| J:\6538\shared\20241016_01_Eric_TCAcells_BEHC18-100\20241016_EMPB__04_10_Std.d | Calibration | 10 | x | 10.0000 | 21438 | 2143.7896 | |
| J:\6538\shared\20241016_01_Eric_TCAcells_BEHC18-100\20241016_EMPB__88_10_Std.d | QC | 10 | x | 10.0000 | 5040 | 503.9518 | 25.1 |
| J:\6538\shared\20241016_01_Eric_TCAcells_BEHC18-100\20241017_EMPB__00_10_Std.d | QC | 10 | x | 10.0000 | 7216 | 721.5529 | 25.1 |
| J:\6538\shared\20241016_01_Eric_TCAcells_BEHC18-100\20241016_EMPB__03_50_Std.d | Calibration | 50 | | 50.0000 | 25400 | 507.9989 | |
| J:\6538\shared\20241016_01_Eric_TCAcells_BEHC18-100\20241016_EMPB__87_50_Std.d | QC | 50 | x | 50.0000 | 10401 | 208.0238 | |
| J:\6538\shared\20241016_01_Eric_TCAcells_BEHC18-100\20241016_EMPB__02_100_Std.d | Calibration | 100 | | 100.0000 | 57204 | 572.0359 | |

### Quantitative Analysis Calibration Report

| | | | |
|---|---|---|---|
| **Batch Data Path** | J:\6538\shared\20241016_01_Eric_TCAcells_BEHC18-100\QuantResults\20241016_EMPB.batch.bin | | |
| **Analysis Time Stamp** | 10/17/2024 2:51:43 PM | **Analyst Name** | MSU\w68p731 |
| **Report Generation Time** | 10/17/2024 3:01:12 PM | **Reporter Name** | MSU\w68p731 |
| **Calibration Last Update** | 10/17/2024 2:44:43 PM | **Batch State** | Processed |
| **Analyze Quant Version** | 12.0 | **Report Quant Version** | 12.0 |

**Fumaric acid** **Relative Standard Error** 63.3

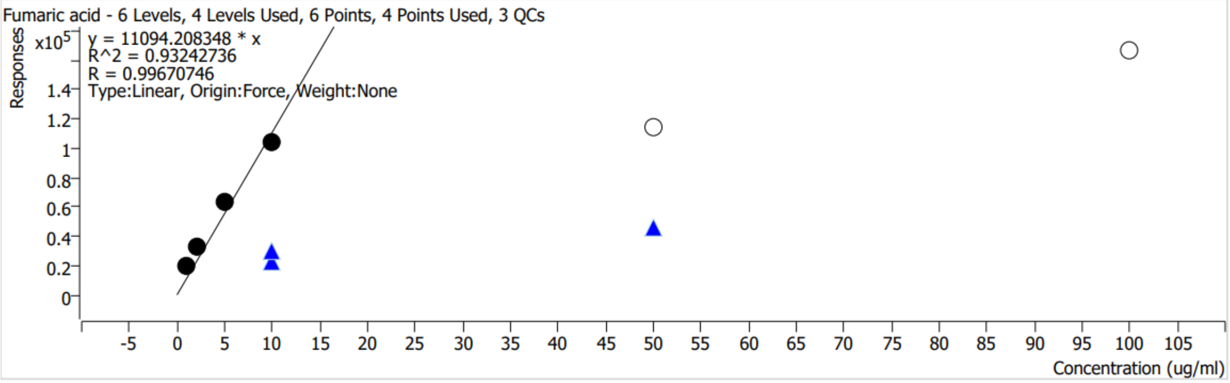


| Calibration STD Path | Type | Level | Enable | Conc. | Response | RF | Level RSD |
|---|---|---|---|---|---|---|---|
| J:\6538\shared\20241016_01_Eric_TCAcells_BEHC18-100\20241016_EMPB__05_1_Std.d | Calibration | 1 | x | 1.0000 | 19313 | 19313.2350 | |
| J:\6538\shared\20241016_01_Eric_TCAcells_BEHC18-100\20241016_EMPB__05_2_Std.d | Calibration | 2 | x | 2.0000 | 32785 | 16392.4565 | |
| J:\6538\shared\20241016_01_Eric_TCAcells_BEHC18-100\20241016_EMPB__05_5_Std.d | Calibration | 5 | x | 5.0000 | 63249 | 12649.7619 | |
| J:\6538\shared\20241016_01_Eric_TCAcells_BEHC18-100\20241016_EMPB__04_10_Std.d | Calibration | 10 | x | 10.0000 | 104112 | 10411.1998 | |
| J:\6538\shared\20241016_01_Eric_TCAcells_BEHC18-100\20241016_EMPB__88_10_Std.d | QC | 10 | x | 10.0000 | 22830 | 2283.0124 | 17.6 |
| J:\6538\shared\20241016_01_Eric_TCAcells_BEHC18-100\20241017_EMPB__00_10_Std.d | QC | 10 | x | 10.0000 | 29339 | 2933.9388 | 17.6 |
| J:\6538\shared\20241016_01_Eric_TCAcells_BEHC18-100\20241016_EMPB__03_50_Std.d | Calibration | 50 | | 50.0000 | 114409 | 2288.1841 | |
| J:\6538\shared\20241016_01_Eric_TCAcells_BEHC18-100\20241016_EMPB__87_50_Std.d | QC | 50 | x | 50.0000 | 46163 | 923.2534 | |
| J:\6538\shared\20241016_01_Eric_TCAcells_BEHC18-100\20241016_EMPB__02_100_Std.d | Calibration | 100 | | 100.0000 | 166094 | 1660.9379 | |

### Quantitative Analysis Calibration Report

| | | | |
|---|---|---|---|
| **Batch Data Path** | J:\6538\shared\20241016_01_Eric_TCAcells_BEHC18-100\QuantResults\20241016_EMPB.batch.bin | | |
| **Analysis Time Stamp** | 10/17/2024 2:51:43 PM | **Analyst Name** | MSU\w68p731 |
| **Report Generation Time** | 10/17/2024 3:01:12 PM | **Reporter Name** | MSU\w68p731 |
| **Calibration Last Update** | 10/17/2024 2:44:43 PM | **Batch State** | Processed |
| **Analyze Quant Version** | 12.0 | **Report Quant Version** | 12.0 |

**Phosphoenol pyruvic acid** **Relative Standard Error** 92.9

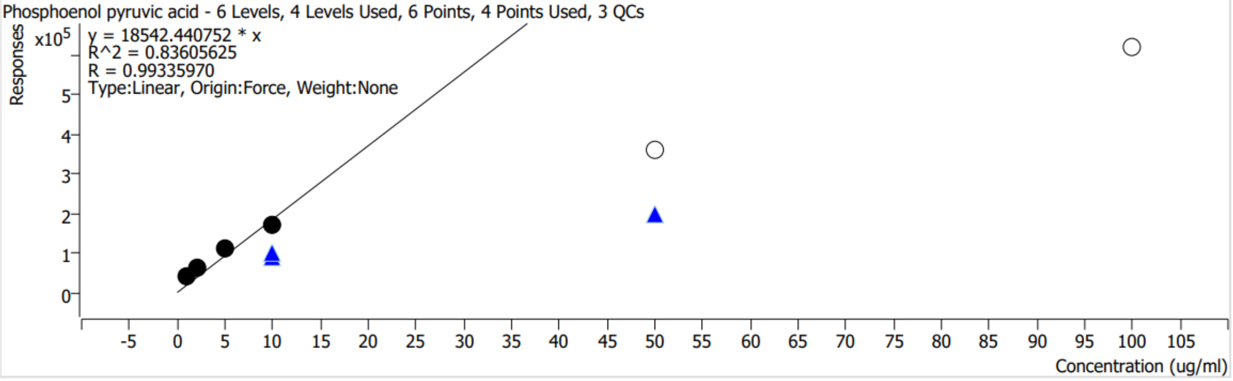


| Calibration STD Path | Type | Level | Enable | Conc. | Response | RF | Level RSD |
|---|---|---|---|---|---|---|---|
| J:\6538\shared\20241016_01_Eric_TCAcells_BEHC18-100\20241016_EMPB__05_1_Std.d | Calibration | 1 | x | 1.0000 | 38922 | 38921.6693 | |
| J:\6538\shared\20241016_01_Eric_TCAcells_BEHC18-100\20241016_EMPB__05_2_Std.d | Calibration | 2 | x | 2.0000 | 62609 | 31304.4547 | |
| J:\6538\shared\20241016_01_Eric_TCAcells_BEHC18-100\20241016_EMPB__05_5_Std.d | Calibration | 5 | x | 5.0000 | 110393 | 22078.5255 | |
| J:\6538\shared\20241016_01_Eric_TCAcells_BEHC18-100\20241016_EMPB__04_10_Std.d | Calibration | 10 | x | 10.0000 | 169441 | 16944.1467 | |
| J:\6538\shared\20241016_01_Eric_TCAcells_BEHC18-100\20241016_EMPB__88_10_Std.d | QC | 10 | x | 10.0000 | 91142 | 9114.2201 | 5.3 |
| J:\6538\shared\20241016_01_Eric_TCAcells_BEHC18-100\20241017_EMPB__00_10_Std.d | QC | 10 | x | 10.0000 | 98246 | 9824.5547 | 5.3 |
| J:\6538\shared\20241016_01_Eric_TCAcells_BEHC18-100\20241016_EMPB__03_50_Std.d | Calibration | 50 | | 50.0000 | 358953 | 7179.0538 | |
| J:\6538\shared\20241016_01_Eric_TCAcells_BEHC18-100\20241016_EMPB__87_50_Std.d | QC | 50 | x | 50.0000 | 195611 | 3912.2131 | |
| J:\6538\shared\20241016_01_Eric_TCAcells_BEHC18-100\20241016_EMPB__02_100_Std.d | Calibration | 100 | | 100.0000 | 618005 | 6180.0474 | |

### Quantitative Analysis Calibration Report

| | | | |
|---|---|---|---|
| **Batch Data Path** | J:\6538\shared\20241016_01_Eric_TCAcells_BEHC18-100\QuantResults\20241016_EMPB.batch.bin | | |
| **Analysis Time Stamp** | 10/17/2024 2:51:43 PM | **Analyst Name** | MSU\w68p731 |
| **Report Generation Time** | 10/17/2024 3:01:12 PM | **Reporter Name** | MSU\w68p731 |
| **Calibration Last Update** | 10/17/2024 2:44:43 PM | **Batch State** | Processed |
| **Analyze Quant Version** | 12.0 | **Report Quant Version** | 12.0 |

**Malic acid** **Relative Standard Error** 702.9

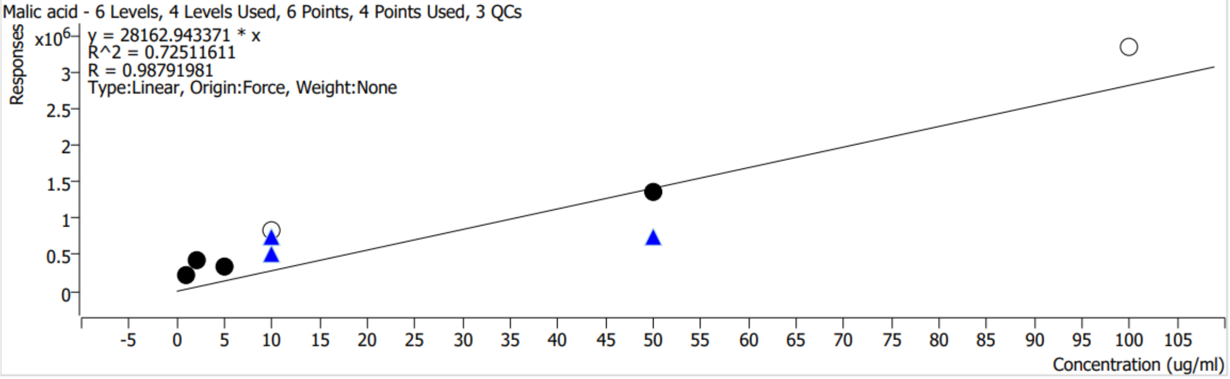


| Calibration STD Path | Type | Level | Enable | Conc. | Response | RF | Level RSD |
|---|---|---|---|---|---|---|---|
| J:\6538\shared\20241016_01_Eric_TCAcells_BEHC18-100\20241016_EMPB__05_1_Std.d | Calibration | 1 | x | 1.0000 | 230497 | 230496.8239 | |
| J:\6538\shared\20241016_01_Eric_TCAcells_BEHC18-100\20241016_EMPB__05_2_Std.d | Calibration | 2 | x | 2.0000 | 434882 | 217441.0116 | |
| J:\6538\shared\20241016_01_Eric_TCAcells_BEHC18-100\20241016_EMPB__05_5_Std.d | Calibration | 5 | x | 5.0000 | 341166 | 68233.1080 | |
| J:\6538\shared\20241016_01_Eric_TCAcells_BEHC18-100\20241016_EMPB__04_10_Std.d | Calibration | 10 | | 10.0000 | 832618 | 83261.8434 | |
| J:\6538\shared\20241016_01_Eric_TCAcells_BEHC18-100\20241016_EMPB__88_10_Std.d | QC | 10 | x | 10.0000 | 515318 | 51531.7608 | 26.6 |
| J:\6538\shared\20241016_01_Eric_TCAcells_BEHC18-100\20241017_EMPB__00_10_Std.d | QC | 10 | x | 10.0000 | 754162 | 75416.1743 | 26.6 |
| J:\6538\shared\20241016_01_Eric_TCAcells_BEHC18-100\20241016_EMPB__03_50_Std.d | Calibration | 50 | x | 50.0000 | 1368923 | 27378.4633 | |
| J:\6538\shared\20241016_01_Eric_TCAcells_BEHC18-100\20241016_EMPB__87_50_Std.d | QC | 50 | x | 50.0000 | 752517 | 15050.3378 | |
| J:\6538\shared\20241016_01_Eric_TCAcells_BEHC18-100\20241016_EMPB__02_100_Std.d | Calibration | 100 | | 100.0000 | 3328353 | 33283.5306 | |

### Quantitative Analysis Calibration Report

| | | | |
|---|---|---|---|
| **Batch Data Path** | J:\6538\shared\20241016_01_Eric_TCAcells_BEHC18-100\QuantResults\20241016_EMPB.batch.bin | | |
| **Analysis Time Stamp** | 10/17/2024 2:51:43 PM | **Analyst Name** | MSU\w68p731 |
| **Report Generation Time** | 10/17/2024 3:01:12 PM | **Reporter Name** | MSU\w68p731 |
| **Calibration Last Update** | 10/17/2024 2:44:43 PM | **Batch State** | Processed |
| **Analyze Quant Version** | 12.0 | **Report Quant Version** | 12.0 |

**2-oxoglutaric acid** **Relative Standard Error** 127.5

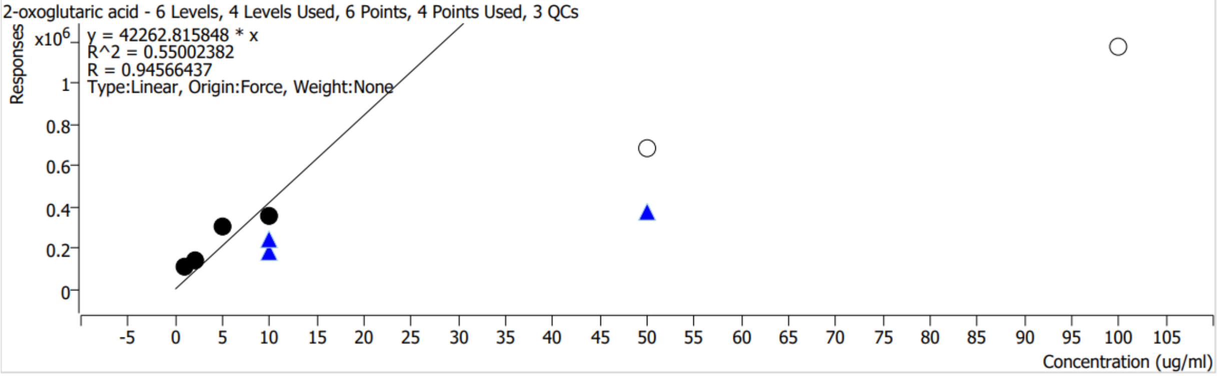


| Calibration STD Path | Type | Level | Enable | Conc. | Response | RF | Level RSD |
|---|---|---|---|---|---|---|---|
| J:\6538\shared\20241016_01_Eric_TCAcells_BEHC18-100\20241016_EMPB__05_1_Std.d | Calibration | 1 | x | 1.0000 | 110469 | 110469.4405 | |
| J:\6538\shared\20241016_01_Eric_TCAcells_BEHC18-100\20241016_EMPB__05_2_Std.d | Calibration | 2 | x | 2.0000 | 140790 | 70395.1849 | |
| J:\6538\shared\20241016_01_Eric_TCAcells_BEHC18-100\20241016_EMPB__05_5_Std.d | Calibration | 5 | x | 5.0000 | 301094 | 60218.8013 | |
| J:\6538\shared\20241016_01_Eric_TCAcells_BEHC18-100\20241016_EMPB__04_10_Std.d | Calibration | 10 | x | 10.0000 | 359665 | 35966.4585 | |
| J:\6538\shared\20241016_01_Eric_TCAcells_BEHC18-100\20241016_EMPB__88_10_Std.d | QC | 10 | x | 10.0000 | 177139 | 17713.9404 | 21.0 |
| J:\6538\shared\20241016_01_Eric_TCAcells_BEHC18-100\20241017_EMPB__00_10_Std.d | QC | 10 | x | 10.0000 | 238920 | 23892.0076 | 21.0 |
| J:\6538\shared\20241016_01_Eric_TCAcells_BEHC18-100\20241016_EMPB__03_50_Std.d | Calibration | 50 | | 50.0000 | 689329 | 13786.5735 | |
| J:\6538\shared\20241016_01_Eric_TCAcells_BEHC18-100\20241016_EMPB__87_50_Std.d | QC | 50 | x | 50.0000 | 372107 | 7442.1371 | |
| J:\6538\shared\20241016_01_Eric_TCAcells_BEHC18-100\20241016_EMPB__02_100_Std.d | Calibration | 100 | | 100.0000 | 1173641 | 11736.4149 | |

### Quantitative Analysis Calibration Report

| | | | |
|---|---|---|---|
| **Batch Data Path** | J:\6538\shared\20241016_01_Eric_TCAcells_BEHC18-100\QuantResults\20241016_EMPB.batch.bin | | |
| **Analysis Time Stamp** | 10/17/2024 2:51:43 PM | **Analyst Name** | MSU\w68p731 |
| **Report Generation Time** | 10/17/2024 3:01:12 PM | **Reporter Name** | MSU\w68p731 |
| **Calibration Last Update** | 10/17/2024 2:44:43 PM | **Batch State** | Processed |
| **Analyze Quant Version** | 12.0 | **Report Quant Version** | 12.0 |

**6-p-gluconate** **Relative Standard Error** 48.7

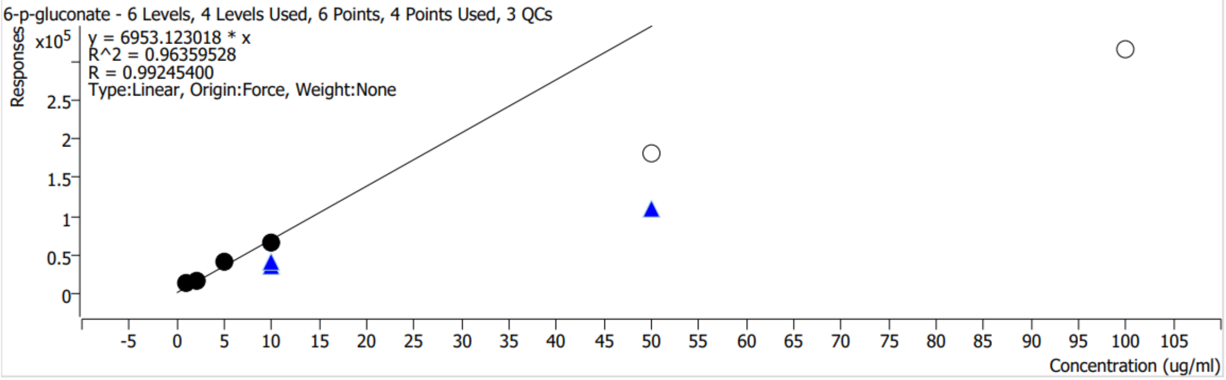


| Calibration STD Path | Type | Level | Enable | Conc. | Response | RF | Level RSD |
|---|---|---|---|---|---|---|---|
| J:\6538\shared\20241016_01_Eric_TCAcells_BEHC18-100\20241016_EMPB__05_1_Std.d | Calibration | 1 | x | 1.0000 | 11551 | 11550.6892 | |
| J:\6538\shared\20241016_01_Eric_TCAcells_BEHC18-100\20241016_EMPB__05_2_Std.d | Calibration | 2 | x | 2.0000 | 14890 | 7444.9646 | |
| J:\6538\shared\20241016_01_Eric_TCAcells_BEHC18-100\20241016_EMPB__05_5_Std.d | Calibration | 5 | x | 5.0000 | 40688 | 8137.6027 | |
| J:\6538\shared\20241016_01_Eric_TCAcells_BEHC18-100\20241016_EMPB__04_10_Std.d | Calibration | 10 | x | 10.0000 | 65914 | 6591.3538 | |
| J:\6538\shared\20241016_01_Eric_TCAcells_BEHC18-100\20241016_EMPB__88_10_Std.d | QC | 10 | x | 10.0000 | 34092 | 3409.1531 | 13.4 |
| J:\6538\shared\20241016_01_Eric_TCAcells_BEHC18-100\20241017_EMPB__00_10_Std.d | QC | 10 | x | 10.0000 | 41224 | 4122.4235 | 13.4 |
| J:\6538\shared\20241016_01_Eric_TCAcells_BEHC18-100\20241016_EMPB__03_50_Std.d | Calibration | 50 | | 50.0000 | 180628 | 3612.5537 | |
| J:\6538\shared\20241016_01_Eric_TCAcells_BEHC18-100\20241016_EMPB__87_50_Std.d | QC | 50 | x | 50.0000 | 109010 | 2180.1951 | |
| J:\6538\shared\20241016_01_Eric_TCAcells_BEHC18-100\20241016_EMPB__02_100_Std.d | Calibration | 100 | | 100.0000 | 316522 | 3165.2152 | |

### Quantitative Analysis Calibration Report

| | | | |
|---|---|---|---|
| **Batch Data Path** | J:\6538\shared\20241016_01_Eric_TCAcells_BEHC18-100\QuantResults\20241016_EMPB.batch.bin | | |
| **Analysis Time Stamp** | 10/17/2024 2:51:43 PM | **Analyst Name** | MSU\w68p731 |
| **Report Generation Time** | 10/17/2024 3:01:12 PM | **Reporter Name** | MSU\w68p731 |
| **Calibration Last Update** | 10/17/2024 2:44:43 PM | **Batch State** | Processed |
| **Analyze Quant Version** | 12.0 | **Report Quant Version** | 12.0 |

**Citric acid** **Relative Standard Error** 141.8

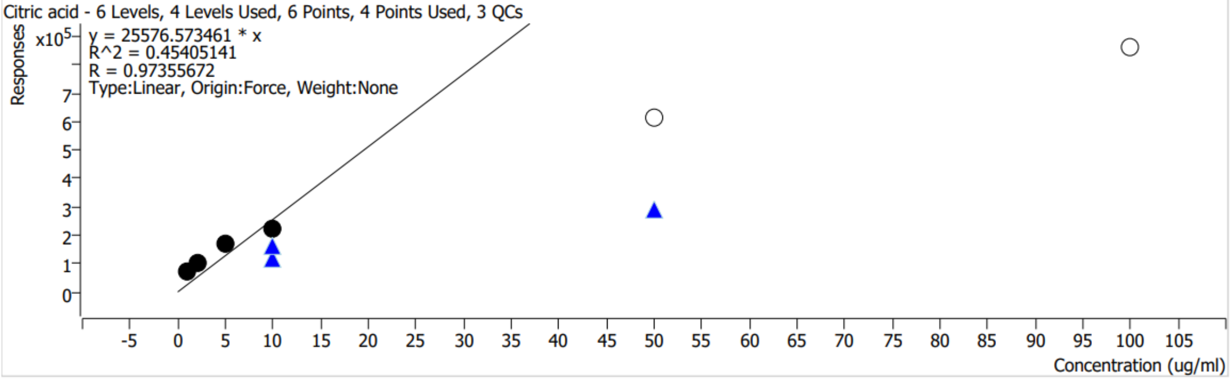


| Calibration STD Path | Type | Level | Enable | Conc. | Response | RF | Level RSD |
|---|---|---|---|---|---|---|---|
| J:\6538\shared\20241016_01_Eric_TCAcells_BEHC18-100\20241016_EMPB__05_1_Std.d | Calibration | 1 | x | 1.0000 | 68773 | 68773.0530 | |
| J:\6538\shared\20241016_01_Eric_TCAcells_BEHC18-100\20241016_EMPB__05_2_Std.d | Calibration | 2 | x | 2.0000 | 103483 | 51741.4495 | |
| J:\6538\shared\20241016_01_Eric_TCAcells_BEHC18-100\20241016_EMPB__05_5_Std.d | Calibration | 5 | x | 5.0000 | 168564 | 33712.7097 | |
| J:\6538\shared\20241016_01_Eric_TCAcells_BEHC18-100\20241016_EMPB__04_10_Std.d | Calibration | 10 | x | 10.0000 | 220640 | 22063.9796 | |
| J:\6538\shared\20241016_01_Eric_TCAcells_BEHC18-100\20241016_EMPB__88_10_Std.d | QC | 10 | x | 10.0000 | 120100 | 12009.9917 | 22.2 |
| J:\6538\shared\20241016_01_Eric_TCAcells_BEHC18-100\20241017_EMPB__00_10_Std.d | QC | 10 | x | 10.0000 | 164786 | 16478.6083 | 22.2 |
| J:\6538\shared\20241016_01_Eric_TCAcells_BEHC18-100\20241016_EMPB__03_50_Std.d | Calibration | 50 | | 50.0000 | 616142 | 12322.8408 | |
| J:\6538\shared\20241016_01_Eric_TCAcells_BEHC18-100\20241016_EMPB__87_50_Std.d | QC | 50 | x | 50.0000 | 287487 | 5749.7379 | |
| J:\6538\shared\20241016_01_Eric_TCAcells_BEHC18-100\20241016_EMPB__02_100_Std.d | Calibration | 100 | | 100.0000 | 858937 | 8589.3657 | |

### Quantitative Analysis Calibration Report

| | | | |
|---|---|---|---|
| **Batch Data Path** | J:\6538\shared\20241016_01_Eric_TCAcells_BEHC18-100\QuantResults\20241016_EMPB.batch.bin | | |
| **Analysis Time Stamp** | 10/17/2024 2:51:43 PM | **Analyst Name** | MSU\w68p731 |
| **Report Generation Time** | 10/17/2024 3:01:12 PM | **Reporter Name** | MSU\w68p731 |
| **Calibration Last Update** | 10/17/2024 2:44:43 PM | **Batch State** | Processed |
| **Analyze Quant Version** | 12.0 | **Report Quant Version** | 12.0 |

**sedoheptulose-7-phosphate** **Relative Standard Error** 131.6

sedoheptulose-7-phosphate - 6 Levels, 5 Levels Used, 6 Points, 5 Points Used, 3 QCs

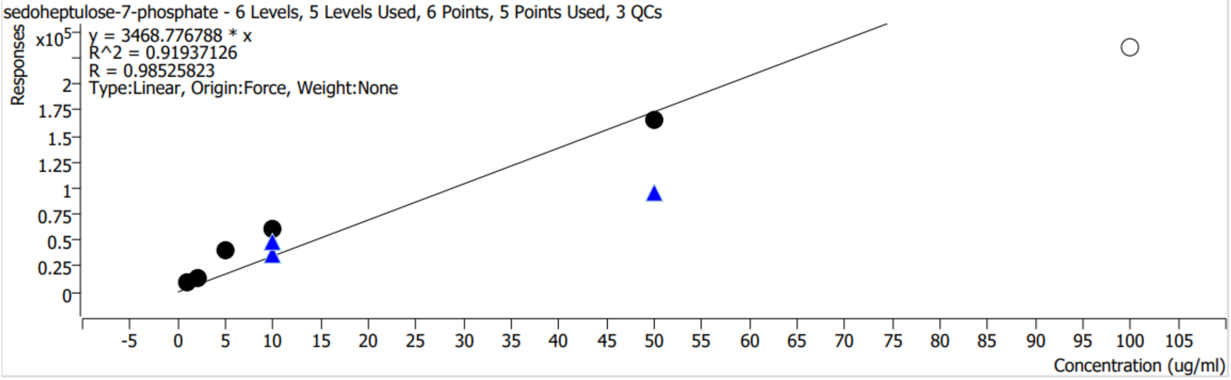


| Calibration STD Path | Type | Level | Enable | Conc. | Response | RF | Level RSD |
|---|---|---|---|---|---|---|---|
| J:\6538\shared\20241016_01_Eric_TCAcells_BEHC18-100\20241016_EMPB__05_1_Std.d | Calibration | 1 | x | 1.0000 | 8513 | 8512.8089 | |
| J:\6538\shared\20241016_01_Eric_TCAcells_BEHC18-100\20241016_EMPB__05_2_Std.d | Calibration | 2 | x | 2.0000 | 13249 | 6624.4038 | |
| J:\6538\shared\20241016_01_Eric_TCAcells_BEHC18-100\20241016_EMPB__05_5_Std.d | Calibration | 5 | x | 5.0000 | 39798 | 7959.5605 | |
| J:\6538\shared\20241016_01_Eric_TCAcells_BEHC18-100\20241016_EMPB__04_10_Std.d | Calibration | 10 | x | 10.0000 | 60929 | 6092.9112 | |
| J:\6538\shared\20241016_01_Eric_TCAcells_BEHC18-100\20241016_EMPB__88_10_Std.d | QC | 10 | x | 10.0000 | 36018 | 3601.8028 | 20.5 |
| J:\6538\shared\20241016_01_Eric_TCAcells_BEHC18-100\20241017_EMPB__00_10_Std.d | QC | 10 | x | 10.0000 | 48245 | 4824.4706 | 20.5 |
| J:\6538\shared\20241016_01_Eric_TCAcells_BEHC18-100\20241016_EMPB__03_50_Std.d | Calibration | 50 | x | 50.0000 | 165592 | 3311.8370 | |
| J:\6538\shared\20241016_01_Eric_TCAcells_BEHC18-100\20241016_EMPB__87_50_Std.d | QC | 50 | x | 50.0000 | 94825 | 1896.4907 | |
| J:\6538\shared\20241016_01_Eric_TCAcells_BEHC18-100\20241016_EMPB__02_100_Std.d | Calibration | 100 | | 100.0000 | 234760 | 2347.6043 | |

### Quantitative Analysis Calibration Report

| | | | |
|---|---|---|---|
| **Batch Data Path** | J:\6538\shared\20241016_01_Eric_TCAcells_BEHC18-100\QuantResults\20241016_EMPB.batch.bin | | |
| **Analysis Time Stamp** | 10/17/2024 2:51:43 PM | **Analyst Name** | MSU\w68p731 |
| **Report Generation Time** | 10/17/2024 3:01:12 PM | **Reporter Name** | MSU\w68p731 |
| **Calibration Last Update** | 10/17/2024 2:44:43 PM | **Batch State** | Processed |
| **Analyze Quant Version** | 12.0 | **Report Quant Version** | 12.0 |

**Succinic acid** **Relative Standard Error** 28.5

Succinic acid - 6 Levels, 6 Levels Used, 6 Points, 6 Points Used, 3 QCs

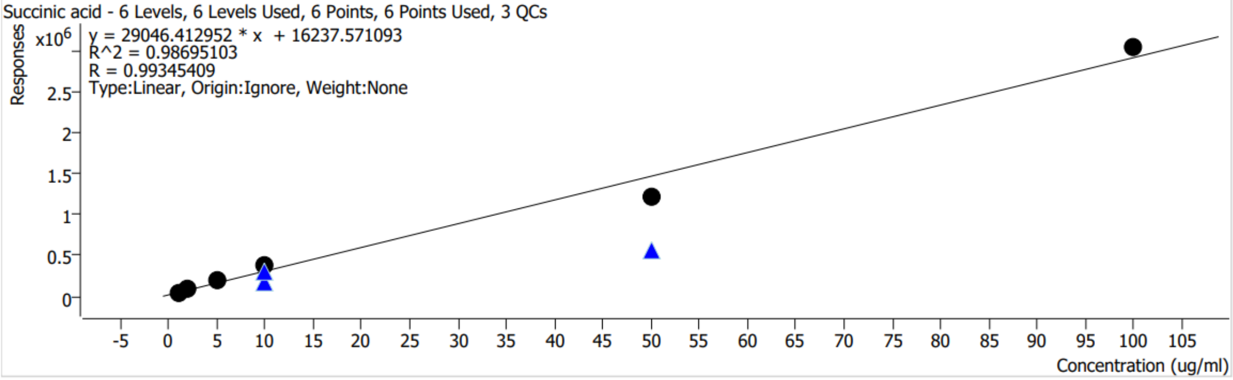


| Calibration STD Path | Type | Level | Enable | Conc. | Response | RF | Level RSD |
|---|---|---|---|---|---|---|---|
| J:\6538\shared\20241016_01_Eric_TCAcells_BEHC18-100\20241016_EMPB__05_1_Std.d | Calibration | 1 | x | 1.0000 | 50746 | 50746.0617 | |
| J:\6538\shared\20241016_01_Eric_TCAcells_BEHC18-100\20241016_EMPB__05_2_Std.d | Calibration | 2 | x | 2.0000 | 95487 | 47743.3427 | |
| J:\6538\shared\20241016_01_Eric_TCAcells_BEHC18-100\20241016_EMPB__05_5_Std.d | Calibration | 5 | x | 5.0000 | 193268 | 38653.6478 | |
| J:\6538\shared\20241016_01_Eric_TCAcells_BEHC18-100\20241016_EMPB__04_10_Std.d | Calibration | 10 | x | 10.0000 | 387204 | 38720.3677 | |
| J:\6538\shared\20241016_01_Eric_TCAcells_BEHC18-100\20241016_EMPB__88_10_Std.d | QC | 10 | x | 10.0000 | 167171 | 16717.1130 | 42.1 |
| J:\6538\shared\20241016_01_Eric_TCAcells_BEHC18-100\20241017_EMPB__00_10_Std.d | QC | 10 | x | 10.0000 | 309095 | 30909.5335 | 42.1 |
| J:\6538\shared\20241016_01_Eric_TCAcells_BEHC18-100\20241016_EMPB__03_50_Std.d | Calibration | 50 | x | 50.0000 | 1210956 | 24219.1215 | |
| J:\6538\shared\20241016_01_Eric_TCAcells_BEHC18-100\20241016_EMPB__87_50_Std.d | QC | 50 | x | 50.0000 | 571050 | 11420.9950 | |
| J:\6538\shared\20241016_01_Eric_TCAcells_BEHC18-100\20241016_EMPB__02_100_Std.d | Calibration | 100 | x | 100.0000 | 3039562 | 30395.6206 | |

### Quantitative Analysis Calibration Report

| | | | |
|---|---|---|---|
| **Batch Data Path** | J:\6538\shared\20241016_01_Eric_TCAcells_BEHC18-100\QuantResults\20241016_EMPB.batch.bin | | |
| **Analysis Time Stamp** | 10/17/2024 2:51:43 PM | **Analyst Name** | MSU\w68p731 |
| **Report Generation Time** | 10/17/2024 3:01:12 PM | **Reporter Name** | MSU\w68p731 |
| **Calibration Last Update** | 10/17/2024 2:44:43 PM | **Batch State** | Processed |
| **Analyze Quant Version** | 12.0 | **Report Quant Version** | 12.0 |

**Fructose-6-phosphate** **Relative Standard Error** 138.0

Fructose-6-phosphate - 6 Levels, 5 Levels Used, 6 Points, 5 Points Used, 3 QCs

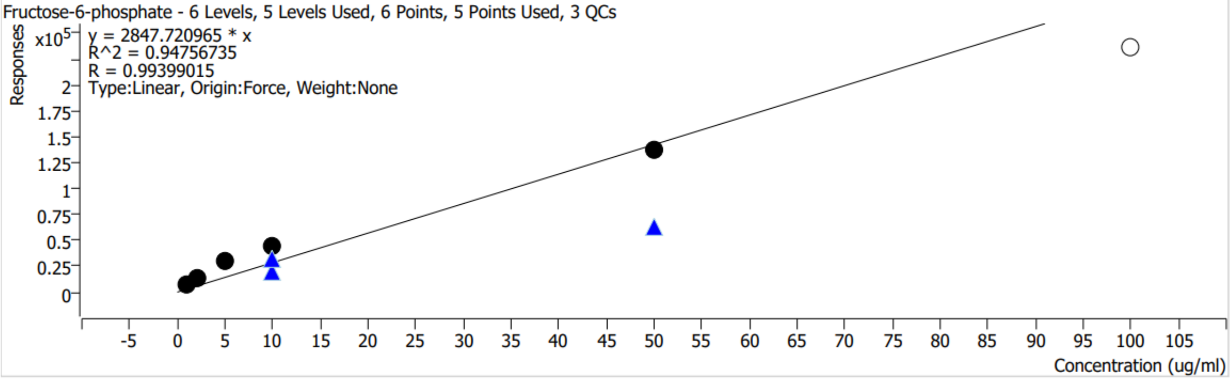


| Calibration STD Path | Type | Level | Enable | Conc. | Response | RF | Level RSD |
|---|---|---|---|---|---|---|---|
| J:\6538\shared\20241016_01_Eric_TCAcells_BEHC18-100\20241016_EMPB__05_1_Std.d | Calibration | 1 | x | 1.0000 | 6752 | 6751.6727 | |
| J:\6538\shared\20241016_01_Eric_TCAcells_BEHC18-100\20241016_EMPB__05_2_Std.d | Calibration | 2 | x | 2.0000 | 14596 | 7298.1885 | |
| J:\6538\shared\20241016_01_Eric_TCAcells_BEHC18-100\20241016_EMPB__05_5_Std.d | Calibration | 5 | x | 5.0000 | 29113 | 5822.5066 | |
| J:\6538\shared\20241016_01_Eric_TCAcells_BEHC18-100\20241016_EMPB__04_10_Std.d | Calibration | 10 | x | 10.0000 | 44094 | 4409.4108 | |
| J:\6538\shared\20241016_01_Eric_TCAcells_BEHC18-100\20241016_EMPB__88_10_Std.d | QC | 10 | x | 10.0000 | 19865 | 1986.5162 | 34.9 |
| J:\6538\shared\20241016_01_Eric_TCAcells_BEHC18-100\20241017_EMPB__00_10_Std.d | QC | 10 | x | 10.0000 | 32874 | 3287.3763 | 34.9 |
| J:\6538\shared\20241016_01_Eric_TCAcells_BEHC18-100\20241016_EMPB__03_50_Std.d | Calibration | 50 | x | 50.0000 | 137341 | 2746.8232 | |
| J:\6538\shared\20241016_01_Eric_TCAcells_BEHC18-100\20241016_EMPB__87_50_Std.d | QC | 50 | x | 50.0000 | 63368 | 1267.3590 | |
| J:\6538\shared\20241016_01_Eric_TCAcells_BEHC18-100\20241016_EMPB__02_100_Std.d | Calibration | 100 | | 100.0000 | 235551 | 2355.5105 | |

### Quantitative Analysis Calibration Report

| | | | |
|---|---|---|---|
| **Batch Data Path** | J:\6538\shared\20241016_01_Eric_TCAcells_BEHC18-100\QuantResults\20241016_EMPB.batch.bin | | |
| **Analysis Time Stamp** | 10/17/2024 2:51:43 PM | **Analyst Name** | MSU\w68p731 |
| **Report Generation Time** | 10/17/2024 3:01:12 PM | **Reporter Name** | MSU\w68p731 |
| **Calibration Last Update** | 10/17/2024 2:44:43 PM | **Batch State** | Processed |
| **Analyze Quant Version** | 12.0 | **Report Quant Version** | 12.0 |

**Glutamic Acid** **Relative Standard Error** 291.6

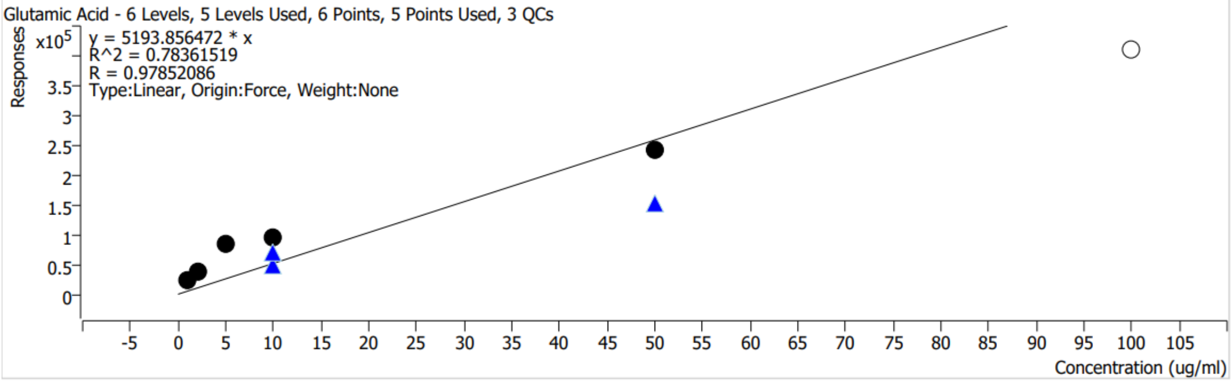


| Calibration STD Path | Type | Level | Enable | Conc. | Response | RF | Level RSD |
|---|---|---|---|---|---|---|---|
| J:\6538\shared\20241016_01_Eric_TCAcells_BEHC18-100\20241016_EMPB__05_1_Std.d | Calibration | 1 | x | 1.0000 | 24178 | 24177.5898 | |
| J:\6538\shared\20241016_01_Eric_TCAcells_BEHC18-100\20241016_EMPB__05_2_Std.d | Calibration | 2 | x | 2.0000 | 36698 | 18349.1413 | |
| J:\6538\shared\20241016_01_Eric_TCAcells_BEHC18-100\20241016_EMPB__05_5_Std.d | Calibration | 5 | x | 5.0000 | 84141 | 16828.2844 | |
| J:\6538\shared\20241016_01_Eric_TCAcells_BEHC18-100\20241016_EMPB__04_10_Std.d | Calibration | 10 | x | 10.0000 | 95877 | 9587.6854 | |
| J:\6538\shared\20241016_01_Eric_TCAcells_BEHC18-100\20241016_EMPB__88_10_Std.d | QC | 10 | x | 10.0000 | 50514 | 5051.4224 | 22.1 |
| J:\6538\shared\20241016_01_Eric_TCAcells_BEHC18-100\20241017_EMPB__00_10_Std.d | QC | 10 | x | 10.0000 | 69193 | 6919.3410 | 22.1 |
| J:\6538\shared\20241016_01_Eric_TCAcells_BEHC18-100\20241016_EMPB__03_50_Std.d | Calibration | 50 | x | 50.0000 | 243656 | 4873.1171 | |
| J:\6538\shared\20241016_01_Eric_TCAcells_BEHC18-100\20241016_EMPB__87_50_Std.d | QC | 50 | x | 50.0000 | 154818 | 3096.3668 | |
| J:\6538\shared\20241016_01_Eric_TCAcells_BEHC18-100\20241016_EMPB__02_100_Std.d | Calibration | 100 | | 100.0000 | 410601 | 4106.0123 | |

### Quantitative Analysis Calibration Report

| | | | |
|---|---|---|---|
| **Batch Data Path** | J:\6538\shared\20241016_01_Eric_TCAcells_BEHC18-100\QuantResults\20241016_EMPB.batch.bin | | |
| **Analysis Time Stamp** | 10/17/2024 2:51:43 PM | **Analyst Name** | MSU\w68p731 |
| **Report Generation Time** | 10/17/2024 3:01:12 PM | **Reporter Name** | MSU\w68p731 |
| **Calibration Last Update** | 10/17/2024 2:44:43 PM | **Batch State** | Processed |
| **Analyze Quant Version** | 12.0 | **Report Quant Version** | 12.0 |

**Lactic acid** **Relative Standard Error** 26.8

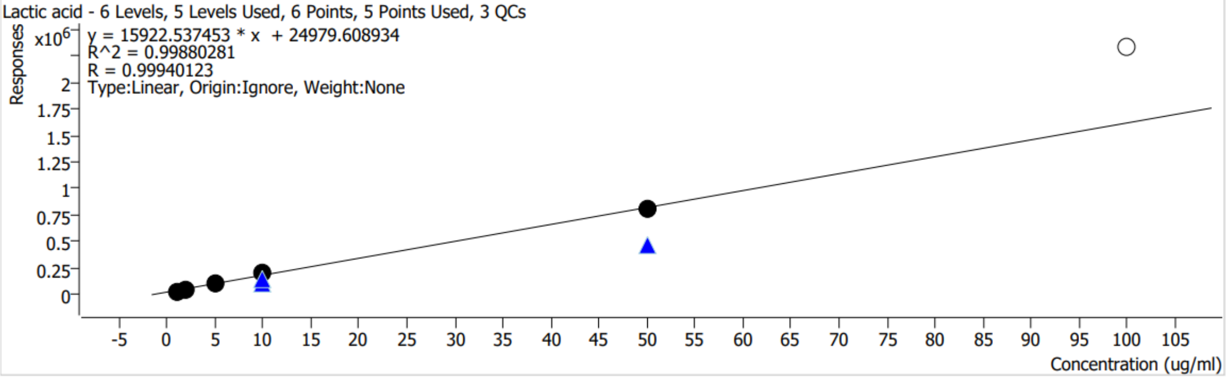


| Calibration STD Path | Type | Level | Enable | Conc. | Response | RF | Level RSD |
|---|---|---|---|---|---|---|---|
| J:\6538\shared\20241016_01_Eric_TCAcells_BEHC18-100\20241016_EMPB__05_1_Std.d | Calibration | 1 | x | 1.0000 | 35047 | 35046.5926 | |
| J:\6538\shared\20241016_01_Eric_TCAcells_BEHC18-100\20241016_EMPB__05_2_Std.d | Calibration | 2 | x | 2.0000 | 48820 | 24410.0634 | |
| J:\6538\shared\20241016_01_Eric_TCAcells_BEHC18-100\20241016_EMPB__05_5_Std.d | Calibration | 5 | x | 5.0000 | 101712 | 20342.3121 | |
| J:\6538\shared\20241016_01_Eric_TCAcells_BEHC18-100\20241016_EMPB__04_10_Std.d | Calibration | 10 | x | 10.0000 | 204224 | 20422.4333 | |
| J:\6538\shared\20241016_01_Eric_TCAcells_BEHC18-100\20241016_EMPB__88_10_Std.d | QC | 10 | x | 10.0000 | 111794 | 11179.4303 | 18.2 |
| J:\6538\shared\20241016_01_Eric_TCAcells_BEHC18-100\20241017_EMPB__00_10_Std.d | QC | 10 | x | 10.0000 | 144844 | 14484.4067 | 18.2 |
| J:\6538\shared\20241016_01_Eric_TCAcells_BEHC18-100\20241016_EMPB__03_50_Std.d | Calibration | 50 | x | 50.0000 | 817828 | 16356.5596 | |
| J:\6538\shared\20241016_01_Eric_TCAcells_BEHC18-100\20241016_EMPB__87_50_Std.d | QC | 50 | x | 50.0000 | 463901 | 9278.0201 | |
| J:\6538\shared\20241016_01_Eric_TCAcells_BEHC18-100\20241016_EMPB__02_100_Std.d | Calibration | 100 | | 100.0000 | 2324781 | 23247.8112 | |

### Quantitative Analysis Calibration Report

| | | | |
|---|---|---|---|
| **Batch Data Path** | J:\6538\shared\20241016_01_Eric_TCAcells_BEHC18-100\QuantResults\20241016_EMPB.batch.bin | | |
| **Analysis Time Stamp** | 10/17/2024 2:51:43 PM | **Analyst Name** | MSU\w68p731 |
| **Report Generation Time** | 10/17/2024 3:01:12 PM | **Reporter Name** | MSU\w68p731 |
| **Calibration Last Update** | 10/17/2024 2:44:43 PM | **Batch State** | Processed |
| **Analyze Quant Version** | 12.0 | **Report Quant Version** | 12.0 |

**Glutamine** **Relative Standard Error** 99.0

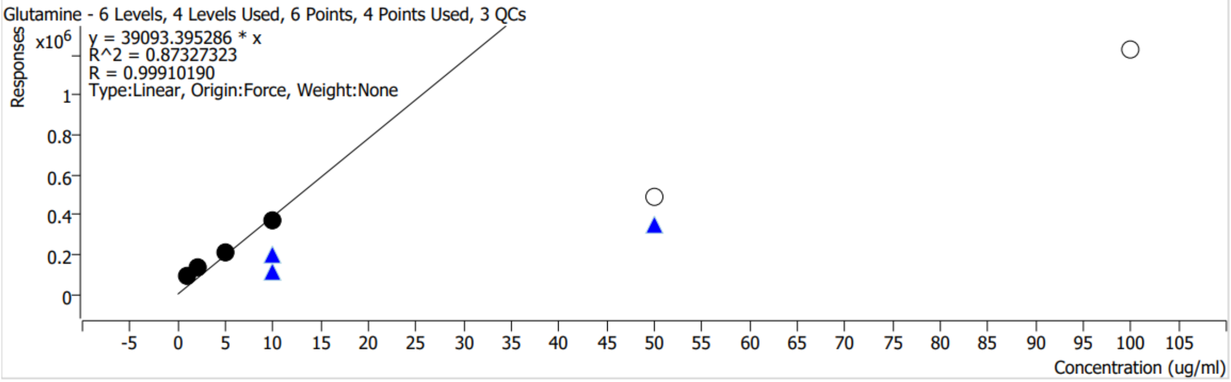


| Calibration STD Path | Type | Level | Enable | Conc. | Response | RF | Level RSD |
|---|---|---|---|---|---|---|---|
| J:\6538\shared\20241016_01_Eric_TCAcells_BEHC18-100\20241016_EMPB__05_1_Std.d | Calibration | 1 | x | 1.0000 | 87134 | 87133.7552 | |
| J:\6538\shared\20241016_01_Eric_TCAcells_BEHC18-100\20241016_EMPB__05_2_Std.d | Calibration | 2 | x | 2.0000 | 129906 | 64953.0716 | |
| J:\6538\shared\20241016_01_Eric_TCAcells_BEHC18-100\20241016_EMPB__05_5_Std.d | Calibration | 5 | x | 5.0000 | 212091 | 42418.2437 | |
| J:\6538\shared\20241016_01_Eric_TCAcells_BEHC18-100\20241016_EMPB__04_10_Std.d | Calibration | 10 | x | 10.0000 | 367474 | 36747.3925 | |
| J:\6538\shared\20241016_01_Eric_TCAcells_BEHC18-100\20241016_EMPB__88_10_Std.d | QC | 10 | x | 10.0000 | 111961 | 11196.0949 | 41.1 |
| J:\6538\shared\20241016_01_Eric_TCAcells_BEHC18-100\20241017_EMPB__00_10_Std.d | QC | 10 | x | 10.0000 | 203651 | 20365.0738 | 41.1 |
| J:\6538\shared\20241016_01_Eric_TCAcells_BEHC18-100\20241016_EMPB__03_50_Std.d | Calibration | 50 | | 50.0000 | 488020 | 9760.4033 | |
| J:\6538\shared\20241016_01_Eric_TCAcells_BEHC18-100\20241016_EMPB__87_50_Std.d | QC | 50 | x | 50.0000 | 350973 | 7019.4623 | |
| J:\6538\shared\20241016_01_Eric_TCAcells_BEHC18-100\20241016_EMPB__02_100_Std.d | Calibration | 100 | | 100.0000 | 1223898 | 12238.9806 | |

### Quantitative Analysis Calibration Report

| | | | |
|---|---|---|---|
| **Batch Data Path** | J:\6538\shared\20241016_01_Eric_TCAcells_BEHC18-100\QuantResults\20241016_EMPB.batch.bin | | |
| **Analysis Time Stamp** | 10/17/2024 2:51:43 PM | **Analyst Name** | MSU\w68p731 |
| **Report Generation Time** | 10/17/2024 3:01:12 PM | **Reporter Name** | MSU\w68p731 |
| **Calibration Last Update** | 10/17/2024 2:44:43 PM | **Batch State** | Processed |
| **Analyze Quant Version** | 12.0 | **Report Quant Version** | 12.0 |

**Fructose/Glucose** **Relative Standard Error** 55.4

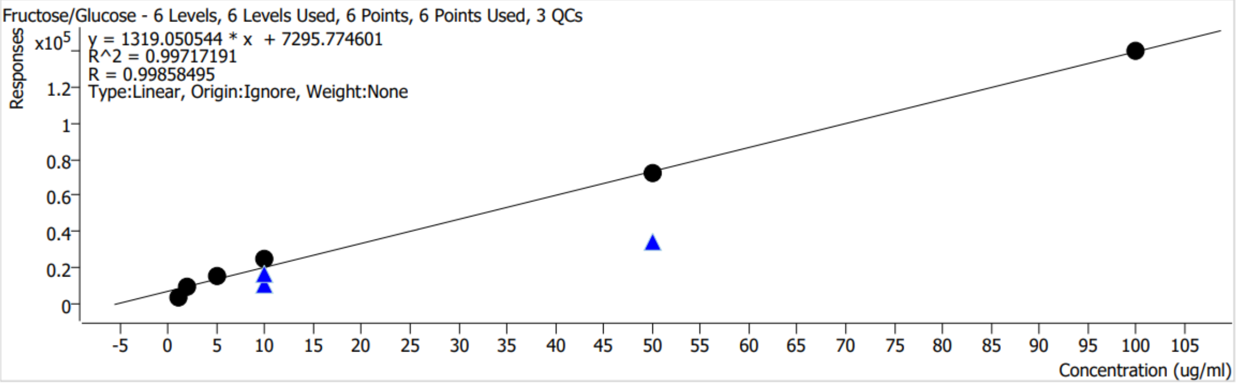


| Calibration STD Path | Type | Level | Enable | Conc. | Response | RF | Level RSD |
|---|---|---|---|---|---|---|---|
| J:\6538\shared\20241016_01_Eric_TCAcells_BEHC18-100\20241016_EMPB__05_1_Std.d | Calibration | 1 | x | 1.0000 | 4373 | 4372.7731 | |
| J:\6538\shared\20241016_01_Eric_TCAcells_BEHC18-100\20241016_EMPB__05_2_Std.d | Calibration | 2 | x | 2.0000 | 9243 | 4621.5065 | |
| J:\6538\shared\20241016_01_Eric_TCAcells_BEHC18-100\20241016_EMPB__05_5_Std.d | Calibration | 5 | x | 5.0000 | 15497 | 3099.4421 | |
| J:\6538\shared\20241016_01_Eric_TCAcells_BEHC18-100\20241016_EMPB__04_10_Std.d | Calibration | 10 | x | 10.0000 | 24658 | 2465.8207 | |
| J:\6538\shared\20241016_01_Eric_TCAcells_BEHC18-100\20241016_EMPB__88_10_Std.d | QC | 10 | x | 10.0000 | 10603 | 1060.2812 | 33.2 |
| J:\6538\shared\20241016_01_Eric_TCAcells_BEHC18-100\20241017_EMPB__00_10_Std.d | QC | 10 | x | 10.0000 | 17117 | 1711.6984 | 33.2 |
| J:\6538\shared\20241016_01_Eric_TCAcells_BEHC18-100\20241016_EMPB__03_50_Std.d | Calibration | 50 | x | 50.0000 | 72440 | 1448.8088 | |
| J:\6538\shared\20241016_01_Eric_TCAcells_BEHC18-100\20241016_EMPB__87_50_Std.d | QC | 50 | x | 50.0000 | 34731 | 694.6155 | |
| J:\6538\shared\20241016_01_Eric_TCAcells_BEHC18-100\20241016_EMPB__02_100_Std.d | Calibration | 100 | x | 100.0000 | 139163 | 1391.6350 | |

### Quantitative Analysis Calibration Report

| | | | |
|---|---|---|---|
| **Batch Data Path** | J:\6538\shared\20241016_01_Eric_TCAcells_BEHC18-100\QuantResults\20241016_EMPB.batch.bin | | |
| **Analysis Time Stamp** | 10/17/2024 2:51:43 PM | **Analyst Name** | MSU\w68p731 |
| **Report Generation Time** | 10/17/2024 3:01:12 PM | **Reporter Name** | MSU\w68p731 |
| **Calibration Last Update** | 10/17/2024 2:44:43 PM | **Batch State** | Processed |
| **Analyze Quant Version** | 12.0 | **Report Quant Version** | 12.0 |

**Pyruvic acid** **Relative Standard Error** 259.0

Pyruvic acid - 6 Levels, 5 Levels Used, 6 Points, 5 Points Used, 3 QCs

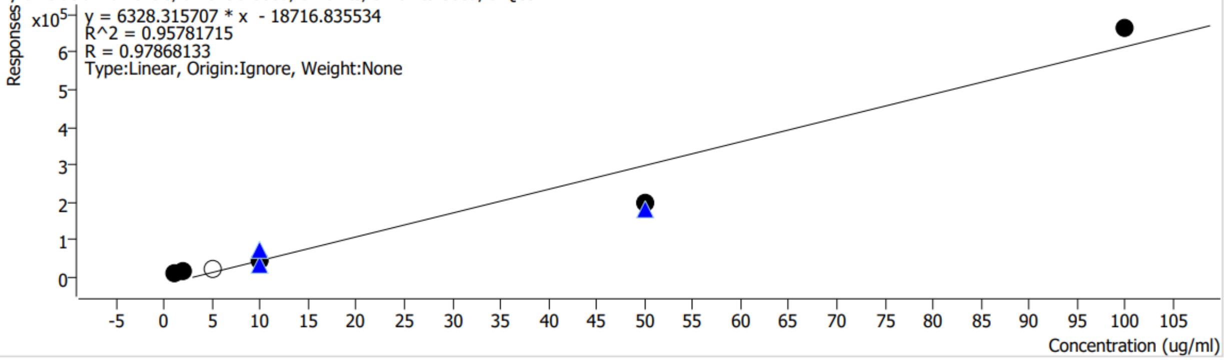


| Calibration STD Path | Type | Level | Enable | Conc. | Response | RF | Level RSD |
|---|---|---|---|---|---|---|---|
| J:\6538\shared\20241016_01_Eric_TCAcells_BEHC18-100\20241016_EMPB__05_1_Std.d | Calibration | 1 | x | 1.0000 | 13440 | 13439.7243 | |
| J:\6538\shared\20241016_01_Eric_TCAcells_BEHC18-100\20241016_EMPB__05_2_Std.d | Calibration | 2 | x | 2.0000 | 17168 | 8583.8982 | |
| J:\6538\shared\20241016_01_Eric_TCAcells_BEHC18-100\20241016_EMPB__05_5_Std.d | Calibration | 5 | | 5.0000 | 20871 | 4174.1839 | |
| J:\6538\shared\20241016_01_Eric_TCAcells_BEHC18-100\20241016_EMPB__04_10_Std.d | Calibration | 10 | x | 10.0000 | 44665 | 4466.4658 | |
| J:\6538\shared\20241016_01_Eric_TCAcells_BEHC18-100\20241016_EMPB__88_10_Std.d | QC | 10 | x | 10.0000 | 36087 | 3608.6610 | 49.5 |
| J:\6538\shared\20241016_01_Eric_TCAcells_BEHC18-100\20241017_EMPB__00_10_Std.d | QC | 10 | x | 10.0000 | 74910 | 7491.0283 | 49.5 |
| J:\6538\shared\20241016_01_Eric_TCAcells_BEHC18-100\20241016_EMPB__03_50_Std.d | Calibration | 50 | x | 50.0000 | 200855 | 4017.1028 | |
| J:\6538\shared\20241016_01_Eric_TCAcells_BEHC18-100\20241016_EMPB__87_50_Std.d | QC | 50 | x | 50.0000 | 182157 | 3643.1356 | |
| J:\6538\shared\20241016_01_Eric_TCAcells_BEHC18-100\20241016_EMPB__02_100_Std.d | Calibration | 100 | x | 100.0000 | 661804 | 6618.0396 | |

### Quantitative Analysis Calibration Report

| | | | |
|---|---|---|---|
| **Batch Data Path** | J:\6538\shared\20241016_01_Eric_TCAcells_BEHC18-100\QuantResults\20241016_EMPB.batch.bin | | |
| **Analysis Time Stamp** | 10/17/2024 2:51:43 PM | **Analyst Name** | MSU\w68p731 |
| **Report Generation Time** | 10/17/2024 3:01:12 PM | **Reporter Name** | MSU\w68p731 |
| **Calibration Last Update** | 10/17/2024 2:44:43 PM | **Batch State** | Processed |
| **Analyze Quant Version** | 12.0 | **Report Quant Version** | 12.0 |

**NADH-1** **Relative Standard Error** 547.3

NADH-1 - 6 Levels, 6 Levels Used, 6 Points, 6 Points Used, 3 QCs

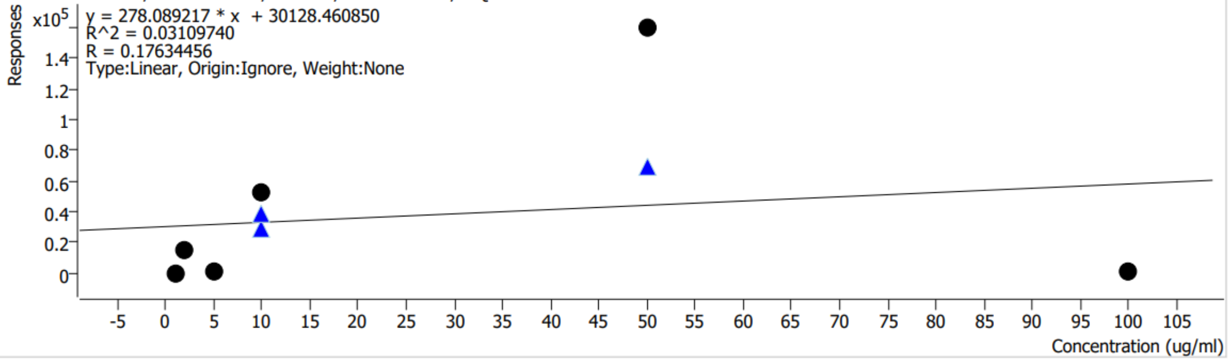


| Calibration STD Path | Type | Level | Enable | Conc. | Response | RF | Level RSD |
|---|---|---|---|---|---|---|---|
| J:\6538\shared\20241016_01_Eric_TCAcells_BEHC18-100\20241016_EMPB__05_1_Std.d | Calibration | 1 | x | 1.0000 | 50 | 50.2955 | |
| J:\6538\shared\20241016_01_Eric_TCAcells_BEHC18-100\20241016_EMPB__05_2_Std.d | Calibration | 2 | x | 2.0000 | 15096 | 7547.9082 | |
| J:\6538\shared\20241016_01_Eric_TCAcells_BEHC18-100\20241016_EMPB__05_5_Std.d | Calibration | 5 | x | 5.0000 | 463 | 92.6655 | |
| J:\6538\shared\20241016_01_Eric_TCAcells_BEHC18-100\20241016_EMPB__04_10_Std.d | Calibration | 10 | x | 10.0000 | 51966 | 5196.6295 | |
| J:\6538\shared\20241016_01_Eric_TCAcells_BEHC18-100\20241016_EMPB__88_10_Std.d | QC | 10 | x | 10.0000 | 29113 | 2911.3293 | 18.8 |
| J:\6538\shared\20241016_01_Eric_TCAcells_BEHC18-100\20241017_EMPB__00_10_Std.d | QC | 10 | x | 10.0000 | 38029 | 3802.9004 | 18.8 |
| J:\6538\shared\20241016_01_Eric_TCAcells_BEHC18-100\20241016_EMPB__03_50_Std.d | Calibration | 50 | x | 50.0000 | 159396 | 3187.9101 | |
| J:\6538\shared\20241016_01_Eric_TCAcells_BEHC18-100\20241016_EMPB__87_50_Std.d | QC | 50 | x | 50.0000 | 69470 | 1389.3914 | |
| J:\6538\shared\20241016_01_Eric_TCAcells_BEHC18-100\20241016_EMPB__02_100_Std.d | Calibration | 100 | x | 100.0000 | 519 | 5.1852 | |

### Quantitative Analysis Calibration Report

| | | | |
|---|---|---|---|
| **Batch Data Path** | J:\6538\shared\20241016_01_Eric_TCAcells_BEHC18-100\QuantResults\20241016_EMPB.batch.bin | | |
| **Analysis Time Stamp** | 10/17/2024 2:51:43 PM | **Analyst Name** | MSU\w68p731 |
| **Report Generation Time** | 10/17/2024 3:01:12 PM | **Reporter Name** | MSU\w68p731 |
| **Calibration Last Update** | 10/17/2024 2:44:43 PM | **Batch State** | Processed |
| **Analyze Quant Version** | 12.0 | **Report Quant Version** | 12.0 |

**HS-CoA** **Relative Standard Error** 89.3

HS-CoA - 6 Levels, 4 Levels Used, 6 Points, 4 Points Used, 3 QCs

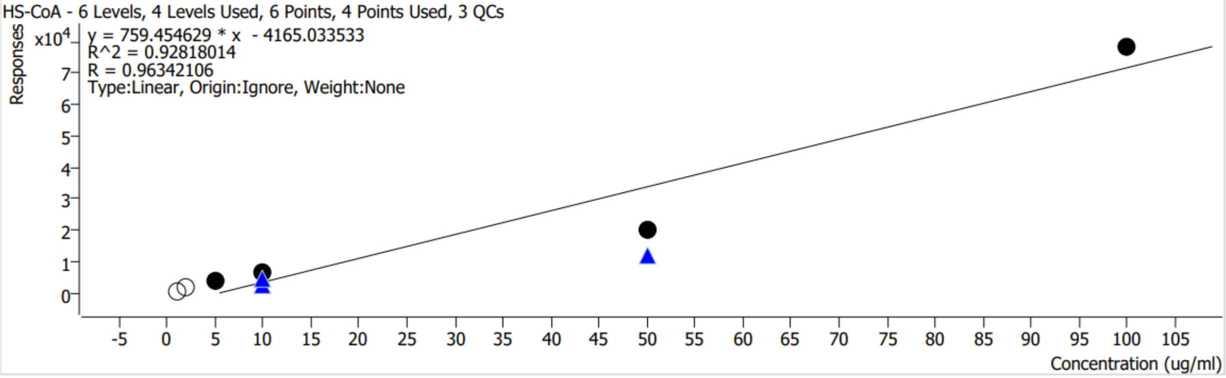


| Calibration STD Path | Type | Level | Enable | Conc. | Response | RF | Level RSD |
|---|---|---|---|---|---|---|---|
| J:\6538\shared\20241016_01_Eric_TCAcells_BEHC18-100\20241016_EMPB__05_1_Std.d | Calibration | 1 | | 1.0000 | 765 | 764.6817 | |
| J:\6538\shared\20241016_01_Eric_TCAcells_BEHC18-100\20241016_EMPB__05_2_Std.d | Calibration | 2 | | 2.0000 | 1921 | 960.6338 | |
| J:\6538\shared\20241016_01_Eric_TCAcells_BEHC18-100\20241016_EMPB__05_5_Std.d | Calibration | 5 | x | 5.0000 | 3937 | 787.4508 | |
| J:\6538\shared\20241016_01_Eric_TCAcells_BEHC18-100\20241016_EMPB__04_10_Std.d | Calibration | 10 | x | 10.0000 | 6557 | 655.7325 | |
| J:\6538\shared\20241016_01_Eric_TCAcells_BEHC18-100\20241016_EMPB__88_10_Std.d | QC | 10 | x | 10.0000 | 2797 | 279.7010 | 29.0 |
| J:\6538\shared\20241016_01_Eric_TCAcells_BEHC18-100\20241017_EMPB__00_10_Std.d | QC | 10 | x | 10.0000 | 4241 | 424.0736 | 29.0 |
| J:\6538\shared\20241016_01_Eric_TCAcells_BEHC18-100\20241016_EMPB__03_50_Std.d | Calibration | 50 | x | 50.0000 | 19998 | 399.9621 | |
| J:\6538\shared\20241016_01_Eric_TCAcells_BEHC18-100\20241016_EMPB__87_50_Std.d | QC | 50 | x | 50.0000 | 12275 | 245.5024 | |
| J:\6538\shared\20241016_01_Eric_TCAcells_BEHC18-100\20241016_EMPB__02_100_Std.d | Calibration | 100 | x | 100.0000 | 78157 | 781.5719 | |

### Quantitative Analysis Calibration Report

Agilent | Trusted Answers

| | | | |
|---|---|---|---|
| **Batch Data Path** | J:\6538\shared\20241016_01_Eric_TCAcells_BEHC18-100\QuantResults\20241016_EMPB.batch.bin | | |
| **Analysis Time Stamp** | 10/17/2024 2:51:43 PM | **Analyst Name** | MSU\w68p731 |
| **Report Generation Time** | 10/17/2024 3:01:12 PM | **Reporter Name** | MSU\w68p731 |
| **Calibration Last Update** | 10/17/2024 2:44:43 PM | **Batch State** | Processed |
| **Analyze Quant Version** | 12.0 | **Report Quant Version** | 12.0 |

**NADH-2** **Relative Standard Error** 206.9

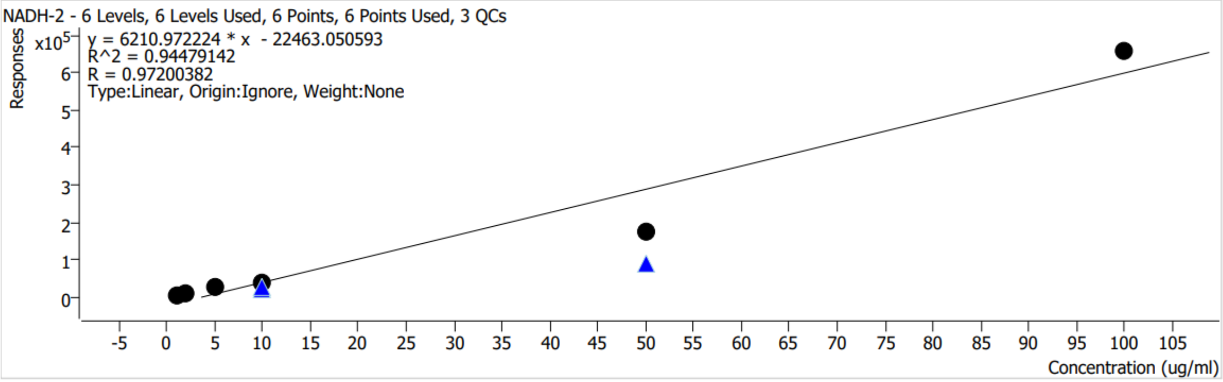


| Calibration STD Path | Type | Level | Enable | Conc. | Response | RF | Level RSD |
|---|---|---|---|---|---|---|---|
| J:\6538\shared\20241016_01_Eric_TCAcells_BEHC18-100\20241016_EMPB__05_1_Std.d | Calibration | 1 | x | 1.0000 | 7219 | 7219.3765 | |
| J:\6538\shared\20241016_01_Eric_TCAcells_BEHC18-100\20241016_EMPB__05_2_Std.d | Calibration | 2 | x | 2.0000 | 8880 | 4440.0325 | |
| J:\6538\shared\20241016_01_Eric_TCAcells_BEHC18-100\20241016_EMPB__05_5_Std.d | Calibration | 5 | x | 5.0000 | 27597 | 5519.4442 | |
| J:\6538\shared\20241016_01_Eric_TCAcells_BEHC18-100\20241016_EMPB__04_10_Std.d | Calibration | 10 | x | 10.0000 | 37466 | 3746.5698 | |
| J:\6538\shared\20241016_01_Eric_TCAcells_BEHC18-100\20241016_EMPB__88_10_Std.d | QC | 10 | x | 10.0000 | 22386 | 2238.6353 | 7.1 |
| J:\6538\shared\20241016_01_Eric_TCAcells_BEHC18-100\20241017_EMPB__00_10_Std.d | QC | 10 | x | 10.0000 | 24742 | 2474.2453 | 7.1 |
| J:\6538\shared\20241016_01_Eric_TCAcells_BEHC18-100\20241016_EMPB__03_50_Std.d | Calibration | 50 | x | 50.0000 | 172342 | 3446.8412 | |
| J:\6538\shared\20241016_01_Eric_TCAcells_BEHC18-100\20241016_EMPB__87_50_Std.d | QC | 50 | x | 50.0000 | 87213 | 1744.2529 | |
| J:\6538\shared\20241016_01_Eric_TCAcells_BEHC18-100\20241016_EMPB__02_100_Std.d | Calibration | 100 | x | 100.0000 | 655161 | 6551.6061 | |
