## Supplemental File 5 for "Targeted analysis of chondrocyte central metabolites in response to cyclical compression and shear deformations"

### Supplemental File 5: Raw Oxygen Data

| Donor Number | Donor Sex | Loading Group | Oxygen Saturation [%] |
|---|---|---|---|
| 3 | F | U | 63.04 |
| | | 15C | 67.91 |
| | | 30C | 64.93 |
| | | 15S | 74.02 |
| | | 30S | 79.10 |
| 4 | M | U | 66.36 |
| | | 15C | 70.98 |
| | | 30C | 76.01 |
| | | 15S | 75.51 |
| | | 30S | 79.68 |
| 5 | F | U | 78.50 |
| | | 15C | 80.05 |
| | | 30C | 78.41 |
| | | 15S | 86.32 |
| | | 30S | 83.72 |
| 6 | M | U | 74.83 |
| | | 15C | 88.54 |
| | | 30C | 82.54 |
| | | 15S | 80.41 |
| | | 30S | 84.32 |
| 7 | F | U | 85.78 |
| | | 15C | 60.37 |
| | | 30C | 53.85 |
| | | 15S | N/A |
| | | 30S | 46.11 |
| 8 | F | U | 74.47 |
| | | 15C | 75.88 |
| | | 30C | 85.28 |
| | | 15S | N/A |
| | | 30S | 83.00 |
| 9 | M | U | 72.11 |
| | | 15C | 74.79 |
| | | 30C | 82.37 |
| | | 15S | 81.21 |
| | | 30S | 74.72 |
| 10 | F | U | 81.60 |
| | | 15C | 86.24 |
| | | 30C | 82.69 |
| | | 15S | 93.24 |
| | | 30S | 76.32 |
